# Restoration of retinal regenerative potential of Müller glia by disrupting intercellular Prox1 transfer

**DOI:** 10.1101/2024.04.23.590652

**Authors:** Eun Jung Lee, Museong Kim, Sooyeon Park, Ji Hyeon Shim, Hyun-Ju Cho, Jung Ah Park, Kihyun Park, Dongeun Lee, Jeong Hwan Kim, Haen Jeong, Fumio Matsuzaki, Seon-Young Kim, Jaehoon Kim, Hanseul Yang, Jeong-Soo Lee, Jin Woo Kim

## Abstract

Individuals with retinal degenerative diseases face significant obstacles in restoring normal vision due to the inability to regenerate degenerated retinal cells. While retinal neurons can be regenerated from Müller glia (MG) in cold-blooded vertebrates, this regenerative ability is lacking in mammals, indicating the limited competence of mammalian MG for retinal regeneration. In this study, we elucidate the role of prospero-related homeobox 1 (Prox1) in rendering the mammalian retina incompetent for MG-derived regeneration. Prox1 accumulates in MG in the degenerating human and mouse retinas but not in those in the regenerating zebrafish retina. Strikingly, Prox1 in mouse MG originated from neighboring retinal neurons through intercellular protein transfer. Consequently, inhibition of Prox1 transfer to MG enables the reprogramming of MG into proliferative retinal progenitor cells in injured mouse retinas. Furthermore, we succeeded in regenerating retinal neurons and delaying vision loss in retinitis pigmentosa disease model in mice by adenoassociated viral gene delivery of an anti-Prox1 antibody that sequesters extracellular Prox1 in the mouse retina. Taken together, our findings highlight the therapeutic potential of anti-Prox1 therapy in restoring the regenerative potential of the mammalian retina.

## Introduction

Cells that constitute animal tissues are born during development and subsequently degenerate during the animal’s lifetime. Degenerated cells can be replaced by new cells in many animal tissues that contain stem cells, such as the skin, liver, and intestines ^1^. Consequently, the functions of these tissues can be maintained despite continuous cell loss. However, neurons in the majority of tissues in the mammalian central nervous system (CNS) cannot be regenerated ^2^, whereas many CNS tissues in cold-blooded vertebrates, such as fishes and amphibians, maintain their regenerative potential ^3,4^. Previous studies have suggested that an absence of promoting factors and/or presence of suppressive factors for a series of regenerative events, including reprogramming, proliferation, and neuronal differentiation of neural stem/progenitor cells (NSCs/NPCs), accounts for the impaired regeneration in mammalian CNS tissues. However, the identities of these regulatory factors remain largely unknown.

Müller glia (MG) in the retina are reprogrammed to NPCs (hereafter MG-derived retinal progenitor cells [MGPCs]) and reenter the cell cycle to regenerate retinal neurons in cold-blooded vertebrates ^3–5^. However, MG cannot be reprogrammed into MGPCs in mammalian retinas, though they partially dedifferentiate before returning to the resting state ^6,7^. One hypothesis explaining the regenerative incompetence of mammalian MG involves the defective expression of key neurogenic factors, such as *achaete-scute family bHLH transcription factor 1* (*Ascl1*) and *lin-28* ^6,8,9^, as evidenced by the ability of ectopically expressed Ascl1 to trigger the reprogramming of MG into MGPCs in the injured mouse retina ^10,11^. However, Ascl1 alone is not sufficient; epigenetic changes, including histone acetylation and DNA demethylation, are also necessary for reprogramming MG into MGPCs ^10–12^.

Prospero-related homeobox 1 (Prox1), an evolutionarily conserved homeobox transcription factor, has been demonstrated to block the proliferation of NSCs/NPCs but induce the differentiation of these cells to the neurons ^13,14^. Furthermore, in a *Drosophila* neuroblast lineage, the Prox1 homolog *prospero* (*pros*) was identified to suppress the expression of *achaete-scute complex* (*asc*) genes ^15^, which are fly *Asc1* homologs. In parallel, *pros* suppresses the expression of *cyclin E* (*CycE*) ^15^ while inducing dacapo (DAP), a cyclin-dependent kinase inhibitor, to suppress post-mitotic neuroblasts from reentering the cell cycle ^16^. These anti-NPC functions are also conserved in mammalian Prox1 ^17–19^.

## Results

### Retinal injury-induced accumulation of Prox1 protein in mouse MG

In developing mouse retina, Prox1 stimulates the differentiation of retinal progenitor cells (RPCs) into horizontal cells (HCs) ^19^. In adult mouse retina, Prox1 is maintained in these retinal neurons, in addition to bipolar cells (BCs) and A2 amacrine cell (AC) subsets. Prox1 could be also detectable in MG at relatively lower level than those in the retinal neurons (Fig. 1a,c) ^20^.

**Figure 1.**
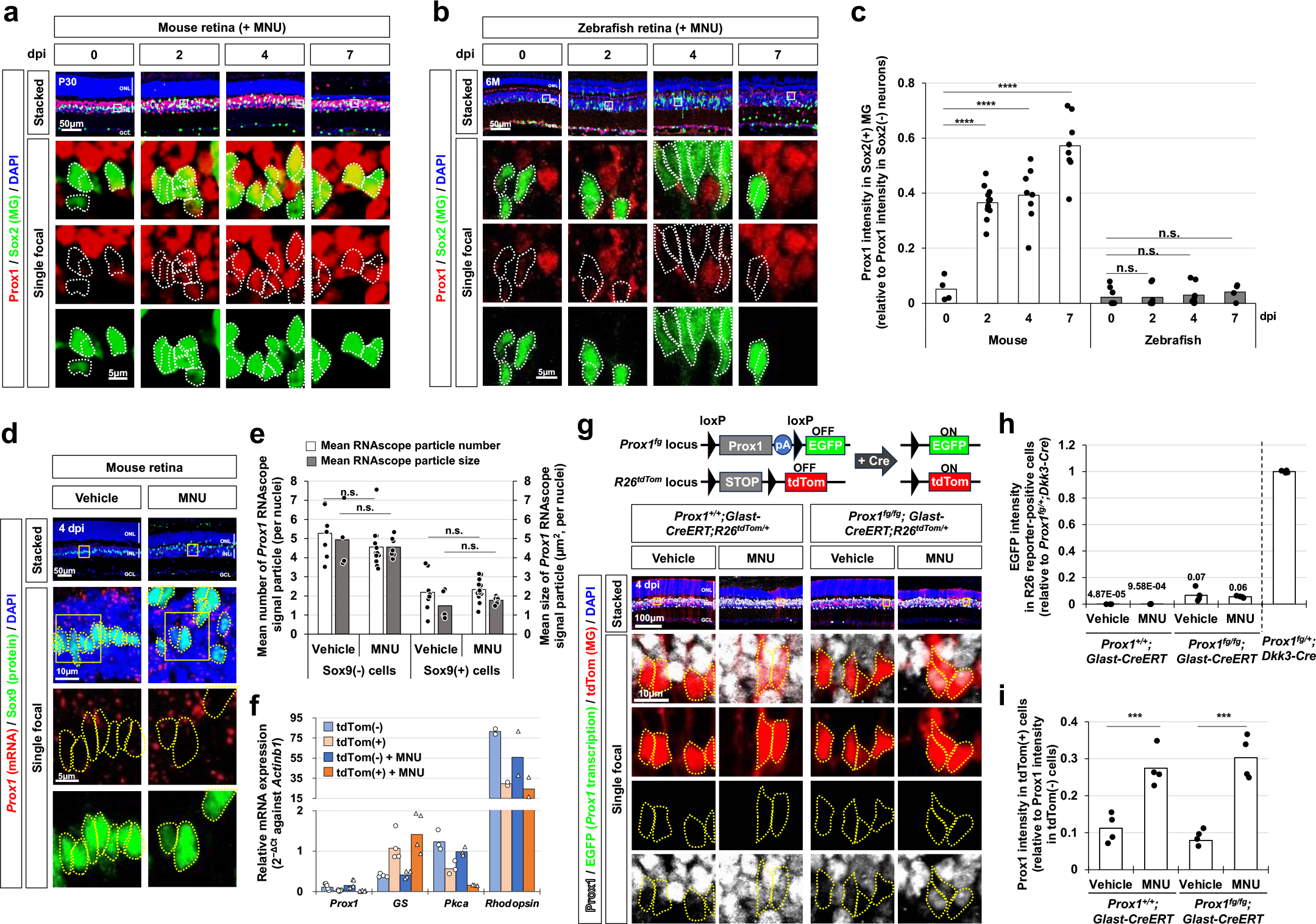
Injury-induced accumulation of exogeneous Prox1 in MG. (**a** and **b**) P30 mice (a) and 6-month (6M)-old zebrafish (b) underwent retinal injury induced by intraperitoneal MNU injection, followed by assessment of Prox1 distribution in the retinas of both species through immunostaining. The single focal images in the bottom rows are magnified versions of the boxed areas in the stacked images in the top row. Sox2-positive MG nuclei are outlined by dotted lines. (**c**) Immunofluorescent intensity of Prox1 in the retinal cells at indicated days post-injury (dpi) was quantified, and the relative Prox1 intensity in MG compared to the retinal neurons in the same image is presented in the graph. Each dot represents the mean intensity collected from a retina. Columns represent mean values (data collected 4 – 6 independent litters). P-values (p) in the graphs were obtained using Student t-tests (* p<0.05; ** p<0.01; *** p<0.005; **** p<0.001; not significant (n.s.), p>0.05). (**d**) *Prox1* mRNA in mouse retinal sections was visualized using RNAscope, in combination with immunostaining of the MG marker, Sox9. (**e**) Mean *Prox1* RNAscope signals in MG nuclei and the rest INL cell nuclei in a retina are presented in the graph (data collected from 3 independent litters). (**f**) tdTom-positive and tdTom-negative retinal cells were isolated from *Glast-CreERT;R26^+/tdTom^* mouse retinas by FACS. Relative mRNA expression of indicated genes to *Actinb1* was determined by their cycle threshold (Ct) scores obtained using RT-qPCR (collected and analyzed from 4 independent batches). (**g**) Complementary EGFP expression from the mouse *Prox1* gene locus upon Cre-dependent excision of *Prox1* cDNA was assessed by co-immunostaining of mouse retinal sections with chicken anti-GFP and rabbit anti- Prox1 antibodies. Cre activity in the same retinal areas was identified by expression of R26^tdTom^ reporter. tdTom-positive cell bodies are outlined by dotted-lines. pA, polyA transcription terminator. (**h**) EGFP intensity of tdTom-positive cells were obtained by confocal microscopy and relative intensity to EGFP intensity of the cells in *Prox1^fg/+^;Dkk3-Cre* mouse retina, provided in Extended Data Fig. 4a, is presented in the graph (data collected from 3 independent litters). The mean scores are also shown in the graph. (**i**) Prox1 intensity in MG relative to that in retinal neurons in the same image is presented in the graph (data collected from 3 independent litters).

Interestingly, Prox1 immunostaining signals in MG were elevated significantly upon N-methyl-N-nitrosourea (MNU)-induced degeneration of photoreceptors (PRs) ^21,22^, whereas Prox1 intensities in other retinal subsets remained unchanged (Fig. 1a,c; Extended Data Fig. 1d). Similar findings were also observed in mouse retinas injured by intensive light exposure that damages PRs; intravitreal injection of N-methyl-D-aspartate (NMDA) that degenerate BCs, ACs, and retinal ganglion cells (RGCs) ^23^; selective degeneration of PRs by diphtheria toxin A (DTA) in *Crx- CreERT2;R26^DTA/+^* mouse retinas treated with an estrogen analog, tamoxifen (Tam) that activates CreERT2 ^24,25^ (Extended Data Fig. 1, a – d). These results suggest that an increase of Prox1 in MG is a common feature of degenerating mouse retinas. In contrast to the results of mouse retinas, the MNU- and NMDA-induced retinal injuries did not increase Prox1 intensity significantly in zebrafish MG, which are known to be reprogrammed to MGPCs in response to the retinal injuries for subsequent proliferation and differentiation to retinal neurons ^3–5^ (Fig. 1b,c; Extended Data Fig. 1e,f).

### Exogenous origin of Prox1 protein in mouse MG

The injury-induced accumulation of Prox1 in MG was an unexpected finding, as the levels of *Prox1* mRNA in MG lineage cells remained consistently low without significant alterations following NMDA- or light-induced injuries to mouse and zebrafish retinas ^6^ (Extended Data Fig. 2). To further explore this phenomenon, we sought to examine changes in *Prox1* mRNA expression in mouse MG using RNAscope *in situ* RNA hybridization. We, however, did not observe an increase of *Prox1* mRNA within the MG following the MNU injury (Fig. 1d,e). This result was further validated through quantitative reverse transcription-polymerase chain reaction (RT-qPCR) analysis of *Prox1* mRNA in tdTomato (tdTom)-positive MG, which were isolated from *Glast-CreERT;R26^tdTom/+^*mouse retinas by fluorescence-activated cell sorting (FACS) (Fig. 1f). In this mouse model, CreERT is selectively expressed in MG by the *glutamate aspartate transporter* (*Glast*) promoter ^26^ and subsequently activated by Tam treatment to express the tdTom Cre reporter, enabling FACS analysis.

Subsequently, we aimed to investigate transcription at the mouse *Prox1* gene locus in MG by utilizing *Prox1^fg^* mice, which are capable of expressing *E<u>G</u>FP* complementarily at the *Prox1* gene locus after the Cre-mediated deletion of the floxed *Prox1* knock-in allele ^18^ (Fig. 1g, diagram). However, the complementary EGFP expression were scarcely observed in tdTom-positive Cre-affected perspective *Prox1*-deleted MG in the Tam-treated *Prox1^fg/fg^;Glast-CreERT;R26^+/tdTom^* mouse retinas (Fig. 1g; Extended Data Fig. 3a). Moreover, the EGFP signals did not show an increase in the tdTom-positive MG following MNU- or NMDA-induced retinal injuries (Fig. 1 g,h; Extended Data Fig. 3a,b). However, Prox1 immunostaining signals were still elevated in these MG cells in the injured *Prox1^fg/fg^;Glast-CreERT;R26^+/tdTom^* mouse retinas (Fig. 1g,i; Extended Data Fig. 3a,c). Taken together, our data suggest that MG in injured mouse retinas accumulate Prox1 without increasing the expression of endogenous *Prox1* gene.

### Exogenous Prox1 suppresses injury-induced proliferation of MG

Previously, we reported that Prox1 protein may have the potential to transfer between cells, a characteristic shared by the majority of homeodomain proteins ^27^. Therefore, it is plausible that mouse MG could uptake Prox1 secreted from neighboring retinal neurons with active *Prox1* gene expression. Our data demonstrate that BCs represent the largest retinal cell population expressing the *Prox1* gene (Extended Data Fig. 4a,b,c). Thus, we investigated whether BCs could serve as a source of Prox1 for MG by genetically eliminating the *Prox1* gene and expressing *EGFP* complementarily in BCs of *Prox1^fg/fg^;Chx10-CreERT2* mice, which express CreERT2 selectively in the BC population in a Tam-dependent manner ^18,28^ (Fig. 2a, top). This manipulation led to a significant reduction in Prox1 intensity in EGFP-negative MG as well as EGFP-positive BCs in *Prox1^fg/fg^;Chx10-CreERT2* mouse retinas compared to those in *Prox1^fg/fg^* littermate mouse retinas, without changes in Prox1 intensity in HCs and ACs (Fig. 2b,d; Extended Data Fig. 5a,b). However, the numbers of BCs or other retinal cells in *Prox1^fg/fg^;Chx10-CreERT2* mice relative to the *Prox1^fg/fg^* littermates did not show significant differences (Extended Data Fig. 5a,c). The visual acuity of the *Prox1^fg/fg^;Chx10-CreERT2* mice was not also significantly different from that of that of the *Prox1^fg/fg^* littermates (Extended Data Fig. 5d). Therefore, these results suggest that the decrease in Prox1 synthesis in BCs, rather than the death or functional impairment of the cells, leads to a reduction in the amount of Prox1 transferred to MG.

**Figure 2.**
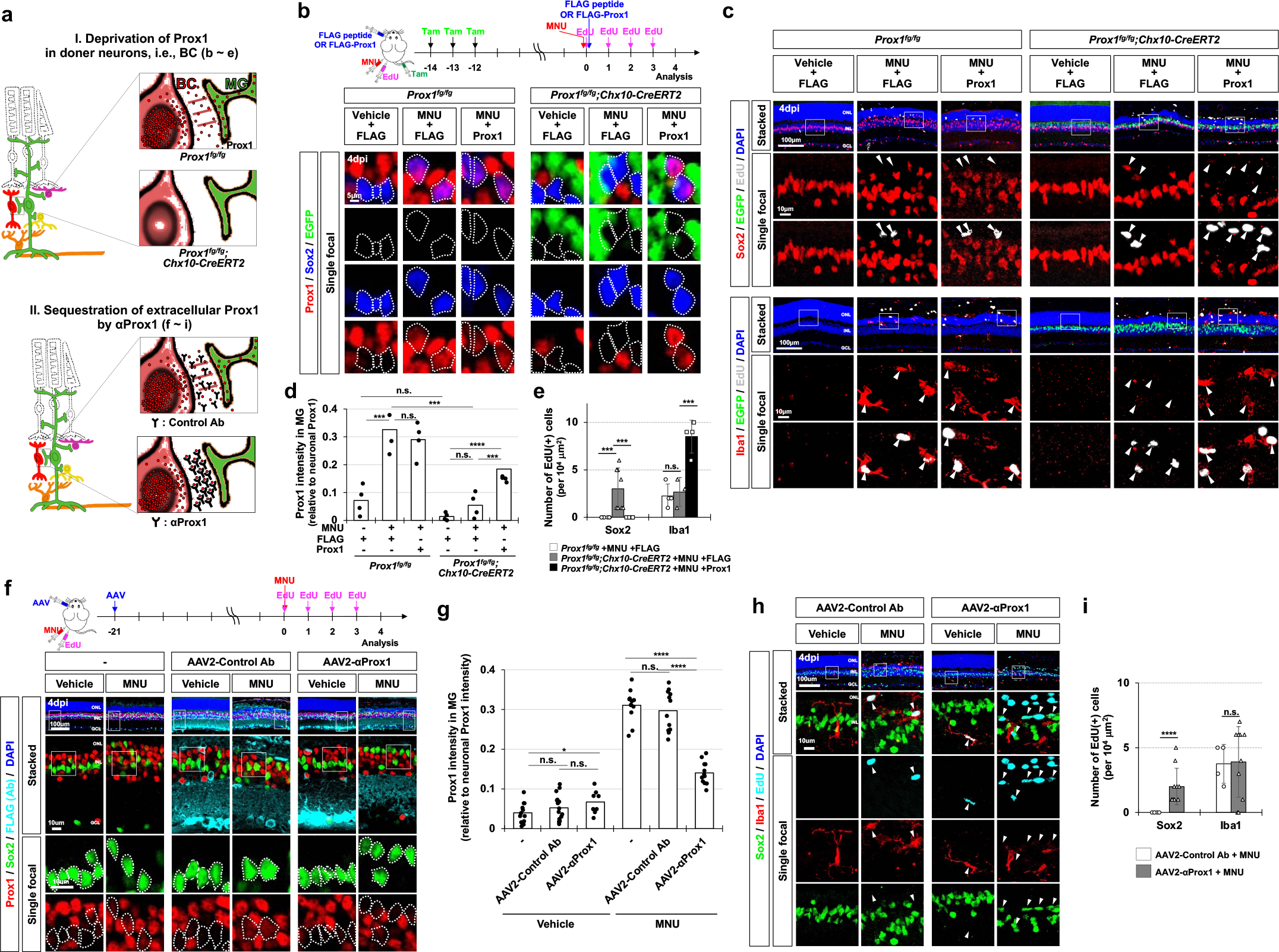
Prox1-depleted MG restore proliferative potential. (**a**) Schematic drawings illustrating the depletion of Prox1 in MG in two approaches: deletion of *Prox1* gene in BC (top) or sequestration of extracellular Prox1 protein by αProx1 (bottom). (**b**) *Prox1* was selectively deleted to express *EGFP* complementarily in BCs of *Prox1^fg/fg^;Chx10-CreERT2* mouse retinas through repeated injections of Tam. The mice were then injected with MNU to degenerate the PRs and EdU to label proliferating cells. 250 fmol FLAG-Prox1 recombinant proteins or FLAG peptides were injected into the vitreal cavity of mouse eyes as indicated. Distribution of Prox1, Sox2, and EGFP in the retinas of *Prox1^fg/fg^* and *Prox1^fg/fg^;Chx10-CreERT2* littermates was examined by immunostaining. Sox2-positive MG nuclei are outlined by dotted-lines. (**c**) MG and microglial identities of EdU-labeled newborn cells in the mouse retinas were determined by co-staining of EdU with Sox2 and Iba1, respectively. Boxed areas in top row are enlarged in next two rows. Arrowheads point EdU-labeled cell nuclei. (**d**) Mean Prox1 intensity in MG relative to that in retinal neurons in the same image is plotted in the graph. (**e**) Numbers of EdU-labeled MG and microglia in the retinas are shown in the graph. (**f**) C57BL/6J mice received intravitreal injections of AAV2- Ctrl Ab or AAV2-αProx1 at a concentration of 5X10^9^ genome copies/eye, followed by injections of MNU and EdU at specified intervals to trace the cells generated post-MNU injury. Retinas of the infected mice were subjected to immunostaining using mouse anti-FLAG antibody to detect FLAG-tagged proteins. Additionally, the distribution of Prox1 in the retinas was also assessed through co-immunostaining with rabbit anti-Prox1 antibody. The MG identity of Prox1-positive cells was determined via co-immunostaining with goat anti-Sox2 antibody. Enlarged views of boxed areas from the top row are presented in the second row, further magnified in the bottom two rows, with Sox2-positive MG nuclei outlined by dotted lines. (**g**) Prox1 intensity in MG relative to that in retinal neurons in the same image is plotted in the graph. (**h**) MG and microglial identities of EdU-labeled cells in the mouse retinas were determined by co-staining of EdU with Sox2 and Iba1, respectively. Boxed areas in top row are magnified in the next rows. Arrowheads point EdU-labeled cell nuclei. (**i**) Numbers of EdU-labeled cells in the retinas are shown in the graph. Graphs in the figure include data collected from 4 independent litters. Error bars denote SEM. * p<0.05; ** p<0.01; ***, p<0.005; **** p<0.001; n.s. >0.05 (ANOVA test).

Remarkably, the number of EdU-labeled proliferative MG significantly increased in the injured retinas of *Prox1^fg/fg^;Chx10-CreERT2* mice (Fig. 2c,e). These findings suggest that MG gain proliferative potential upon the decrease of exogenous Prox1 from retinal neurons. To test this hypothesis, we supplied Prox1 to the retina by intravitreal injection of FLAG-tagged recombinant Prox1 protein to *Prox1^fg/fg^;Chx10-CreERT2* mouse eyes (Fig. 2b, diagram). The FLAG-Prox1 was detectable in MNU-injured mouse MG and its presence was sustained for several days (Extended Data Fig. 6). However, it failed to penetrate vehicle-treated uninjured mouse retinas, which maintained an intact inner retina-blood barrier (Extended Data Fig. 6). The delivery of FLAG- Prox1 resulted in a decrease in the number of EdU-labeled MG but an elevation in the number of EdU-labeled microglia in the injured retinas of *Prox1^fg/fg^;Chx10-CreERT2* mice (Fig. 2c,e).

To further investigate the inverse relationship between the exogenous Prox1 protein accumulation and MG cell proliferation, we also injected FLAG-Prox1 into zebrafish eyes (Extended Data Fig. 7a), which express *glial fibrillary acidic protein* (*gfap*)*-EGFP* transgene in their MG ^29^. The FLAG-Prox1 was detectable in EGFP-positive MG of MNU-injured *gfap:EGFP* zebrafish retinas (Extended Data Fig. 7b,c). This manipulation led to a decrease in the number of Ascl1a-positive MGPCs in these fish retinas (Extended Data Fig. 7d,e). It also reduced the EdU-positivity of Sox2;EGFP-positive MG or MGPC cells (Extended Data Fig. 7, f, k, and l).

Consequently, the regeneration of retinal neurons, identified by their incorporation of EdU and loss of MG-specific gfap-EGFP expression but gain of cell type-specific marker expression, was diminished in the injured zebrafish retinas injected with FLAG-Prox1 (Extended Data Fig. 7, g – l).

### Recovery of MG proliferation in injured mouse retina by sequestering extracellular Prox1

Alternatively, we also aimed to disrupt Prox1 transfer to MG by sequestering it in the extracellular space using an anti-Prox1 antibody (Fig. 2a, bottom). We prepared a cDNA encoding a single-chain fragment (scFv) of chicken immunoglobulin with an affinity for Prox1, which effectively blocks the transfer of Prox1 between cultured cells when added to the growth medium (Extended Data Fig. 8). The anti-Prox1 scFv antibody (αProx1) was fused with the signal sequence of human interleukin-2 (IL-2) before introduction into the genome of adeno-associated virus (AAV), which is known for its effective gene delivery to the mouse retina ^30^. The resulting AAV2/2 serotype viruses (AAV2-αProx1) were injected into the vitreal cavity of mouse eyes to infect retinal cells with the viruses and secrete αProx1 into the extracellular space of the retina (Fig. 2f, diagram). We observed that the accumulation of Prox1 in the MG of MNU-injured mouse retinas was impeded by AAV2-αProx1 infection, whereas infection with AAV2-Control antibody (AAV2-Ctrl Ab), expressing an scFv with no affinity to Prox1, had no such effect (Fig. 2f,g). Furthermore, we noted that the number of MG incorporating EdU for 4 days post-MNU injury was elevated in mouse retinas infected with AAV2-αProx1 compared to those infected with AAV2-Ctrl Ab (Fig. 2h,i).

Taken together, our findings suggest that Prox1 originated from neighboring retinal neurons, including BCs, exerts a suppressive role in injury-induced proliferation of MG and/or transition to proliferative MGPCs through intercellular transfer.

### Transition of Prox1-depleted MG into proliferative MGPCs in the injured mouse retina

Next, we investigated whether the transition of MG to proliferative MGPCs really occurred in MNU-injured *Prox1^fg/fg^;Chx10-CreERT2* mouse retinas through single-cell RNA sequencing (scRNA-seq) analysis (Fig. 3, a – f; Extended Data Fig. 9 – 12; see details in Methods). MG populations in MNU-injured mouse retinas could be categorized into three clusters (Extended Data Fig. 10a). Cluster #3 was detected in both healthy and injured mouse retinas, indicating it represents the resting MG population (Extended Data Fig. 10, b – g). Conversely, clusters #1 and #7, representing injury-induced activated MG populations, were selectively enriched in the injured mouse retinas (Extended Data Fig. 10, b – g). This was supported by the cell lineage flow from cluster #3 to cluster #7 in the velocity plot of scRNA-seq data (Extended Data Fig. 11a).

**Figure 3.**
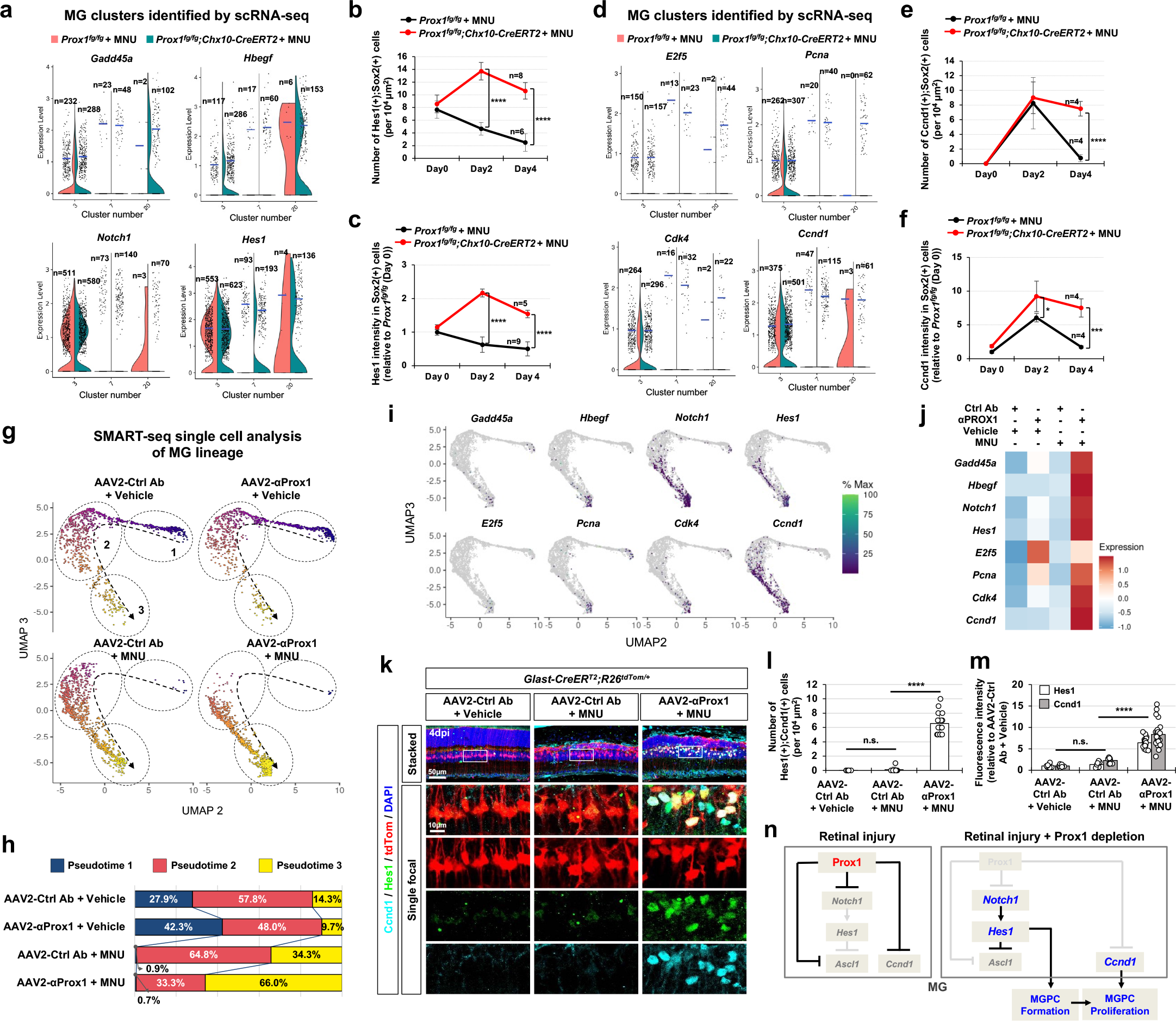
Injury-induced reprogramming of MG into proliferative MGPCs upon decrease in Prox1 transfer. (**a**) Violin plots depicting the expression levels of *Gadd45a*, *Hbegf*, *Notch1*, and *Hes1* mRNA in the indicated cell clusters identified through scRNA-seq analyses of MNU-injured *Prox1^fg/fg^* and *Prox1^fg/fg^;Chx10-CreERT2* mice (details in Extended Data Fig. 9 and 10). Dots represent gene expression levels in individual cells, with the number (n) of dots provided in each column. Blue horizontal bars indicate mean expression values. (**b** and **c**) Distribution of Hes1 in MNU-injured *Prox1^fg/fg^* and *Prox1^fg/fg^;Chx10-CreERT2* mice mouse retinas was examined by immunostaining (images provided in Extended Data Fig. 11c), and the numbers of Hes1- expressing MG (b) and the intensity of Hes1 in MG (c) are shown in the graphs. The numbers of retinas analyzed are provided in the graphs. Error bars denote SEM. (**d**) Violin plots illustrating the expression levels of *E2f5*, *Pcna*, *Cdk4*, and *Ccnd1* mRNA in the indicated cell clusters. (**e** and **f**) Distribution of Ccnd1 in mouse retinas was examined by immunostaining (images provided in Extended Data Fig. 11e), and the numbers of Ccnd1-expressing MG (e) and the intensity of Ccnd1 in MG (f) are shown in the graphs. (**g**) Uniform Manifold Approximation and Projection (UMAP) plots obtained by Monocle 3 analysis of SMART-seq data from FACS purified tdTom- positive cells in Vehicle- or MNU-treated *Glast-CreERT;R26^tdTom/+^*mouse retinas, which were infected with AAV2-Ctrl Ab or AAV2-αProx1 (details in Methods). Dotted arrows delineate the direction of pseudotime progression, while circled areas denote distinct populations identified by numbered pseudotimes. (**h**) Pseudotime distribution of cell populations in each sample is plotted. (**i**) Pseudotime distribution of cells expressing the indicated mRNA of MGPC and proliferating cell markers. (**j**) Mean expression levels of indicated mRNA in each sample displayed in the heatmap. (**k**) Distribution of Ccnd1 and Hes1 in AAV2-infected *Glast-CreERT;R26^tdTom/+^* mouse retinas examined by immunostaining. Boxed areas in top row are magnified in the next rows. (**l** and **m**) The numbers of Ccnd1;Hes1 double-positive cells (l) and the relative intensity of Ccnd1 and Hes1 in those cells (m) are shown in the graphs (data collected from 3 independent litters). (**n**) Hypothetical model illustrating the negative regulation of injury-induced reprogramming of MG to MGPCs by Prox1.

We also observed an additional lineage flow to cluster #7 from cluster #20 (Extended Data Fig. 11a), which was detected at significant cell numbers only in the injured *Prox1^fg/fg^;Chx10- CreERT2* mouse retinas (Extended Data Fig. 10, b – g). Cluster #20 also exhibited lineage flows to the BC (#6) and AC (#14) clusters, suggesting a potential progenitor cell identity for this cluster (Extended Data Fig. 11a). This was supported by the expression of the RPC markers, *growth arrest and DNA damage alpha and gamma* (*Gadd45a and Gadd45g*) ^31^, in this cluster (Fig. 3a; Extended Data Fig. 11b and 12a). The cluster also showed the expression of *heparin-binding epidermal growth factor* (*Hbegf*) and its downstream *early growth response protein 1* (*Egr1*), which induce reprogramming of MG to MGPC ^32,33^, although they did not contain *Ascl1* (Fig. 3a; Extended Data Fig. 11b, 12a, and 12b). Moreover, the cells expressing *hairy and enhancer of split-1* (*Hes1*) and its upstream *Notch1*, which are necessary for the dedifferentiation/proliferation of MG and the maintenance of the dedifferentiated MG ^32–34^, were identified in cluster #20 (Fig. 3a; Extended Data Fig. 11b). Notably, the Hes1 level peaked at 2 days post-MNU injury in Sox2-positive MG or MGPCs in the retinas of *Prox1^fg/fg^;Chx10-CreERT2* mice, while it decreased in the retinas of *Prox1^fg/fg^* mice (Fig. 3b,c; Extended Data Fig. 11c).

The cells in the cluster #20 also expressed *E2F transcription factor 5* (*E2f5*), *Myc*, and *proliferating cell nuclear antigen* (*Pcna*) (Fig. 3d; Extended Data Fig. 11d and 12c), which are enriched in proliferative MGPCs ^6^. Additionally, *Cyclin d1* (*Ccnd1*) and *e1* (*Ccne1*), negative transcription targets of Prox1 in NSCs ^15,16,35^, along with *Cdk4*, which forms a complex with Ccnd1 to trigger G1-to-S cell cycle progression ^36,37^, were detected in cluster #20 cells (Fig. 3d; Extended Data Fig. 11d and 12d). The presence of Ccnd1-expressing MG or MGPCs was elevated and sustained in the MNU-injured retinas of *Prox1^fg/fg^;Chx10-CreERT2* mice, whereas in the retinas of *Prox1^fg/fg^* mice, they appeared transiently and disappeared (Fig. 3e,f; Extended Data Fig. 11e).

Together, these results suggest that cluster #20 represents proliferative MGPCs.

We further investigated injury-induced gene expression changes selectively in MG cell lineages. The tdTom-labeled MG and their descendant cells were isolated via FACS from uninjured or MNU-injured *Glast-CreERT;R26td^Tom/+^* mouse retinas, which were infected with either AAV2-Ctrl Ab or AAV2-αProx1, and mRNA expression patterns of individual cells were analyzed using SMART (Switching Mechanism At the 5’ end of RNA Template) sequencing methods (see details in Methods). We could identify a lineage flow from resting MG to RPC-like cells via activated MG, along with pseudotime progression (Fig. 3g,h). Proliferating cell markers *E2f5*, *Pcna*, *Cdk4*, and *Ccnd1*, as well as MGPC markers, including *Gadd45a, Hbegf*, *Notch1*, and *Hes1*, were detectable in cells at late pseudotime, with these cells being more enriched in MNU- injured mouse retinas infected with AAV2-αProx1 than in other groups (Fig. 3i,j). Consistently, Hes1 and Ccnd1 were induced at significant levels in tdTom-positive MG lineage cells of AAV2- αProx1-infected mouse retina upon MNU injury (Fig. 3k,l,m). Collectively, our results suggest that exogenous Prox1 blocks the conversion of MG to proliferative MGPCs by suppressing the expression of target genes such as *Notch1* and *Ccnd1* ^15,17^. Therefore, inhibiting the transfer of Prox1 to MG is necessary for reprogramming to proliferative MGPCs, consequently the retina could recover the regenerative potential upon retinal injuries (Fig. 3n).

### Delaying vision loss in early onset retinitis pigmentosa mouse models by blocking Prox1 transfer

Despite the development of various therapeutic interventions for retinal degenerative diseases, the majority of patients cannot recover their vision as their lost retinal cells are not regenerated. Therefore, the blocking PROX1 transfer could be a potent therapeutic strategy for retinal degenerative diseases, if exogenous PROX1 also play the suppressive roles in human retinal regeneration. Indeed, PROX1 was accumulated in SOX2-positive MG of 79 years-old retinitis pigmentosa (RP) patient retina but not in the MG of 83 years-old healthy donor retina (Fig. 4a,c). The elevation of Prox1 was also observed in the MG of *Pde6b^rd^*^10^*^/rd^*^10^ mice (Fig. 4d,e), which carry retinal dystrophy-10 (*rd10*) mutations in the phosphodiesterase 6b gene (*Pde6b*) and lose their vision due to the degeneration of rod photoreceptors (rPRs), followed by the loss of cone photoreceptors (cPRs) ^38,39^.

**Figure 4.**
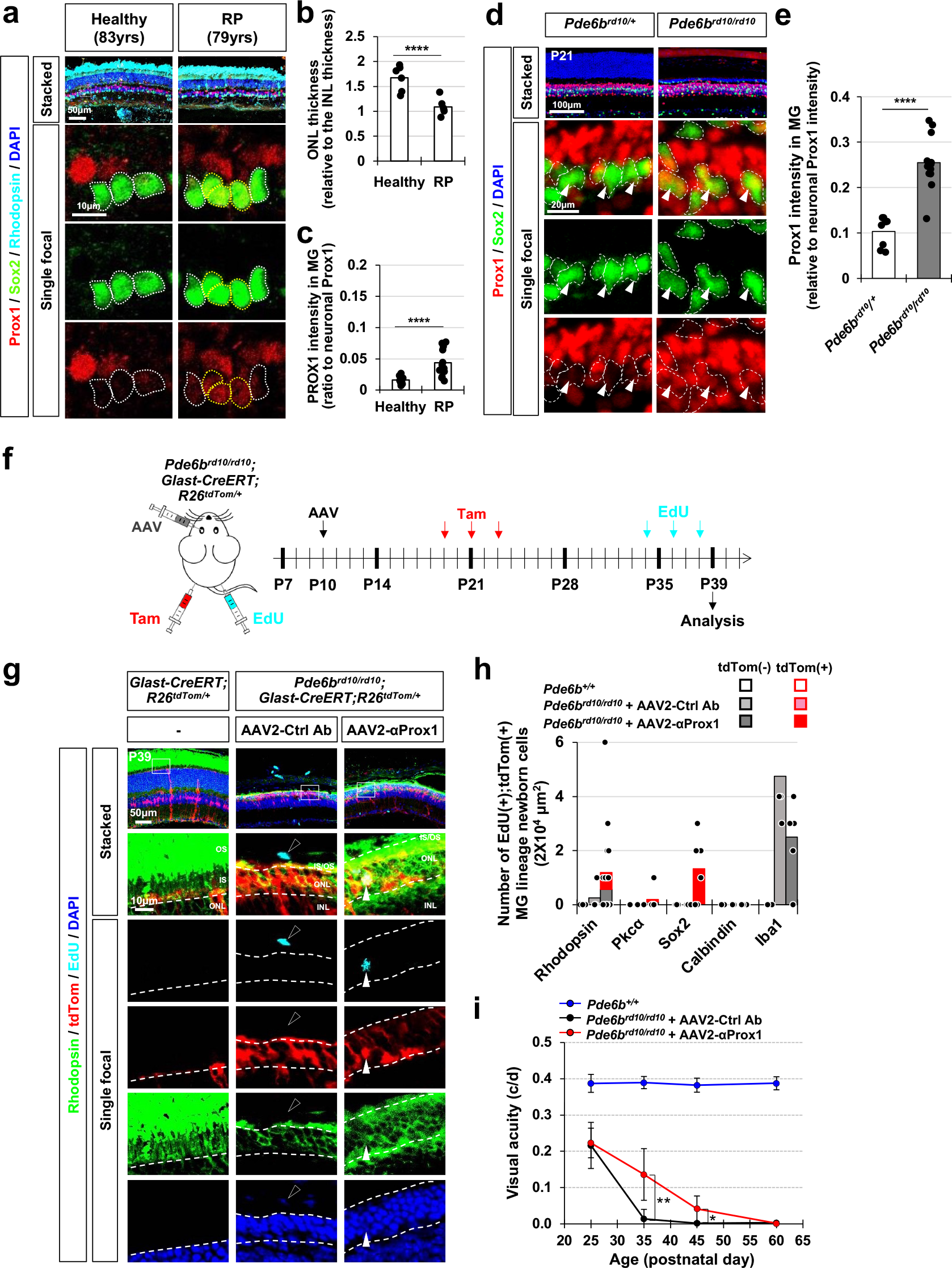
Delaying vision loss in early onset RP model mice by viral gene delivery of anti- Prox1 antibody. (a) Retinal sections from a healthy 83-year-old donor eye and a 79-year-old RP patient eye were stained with rabbit anti-Prox1, goat anti-Sox2, and mouse anti-Rhodopsin antibodies. Nuclei of the cells in the retinal sections were visualized by DAPI staining. Sox2- positive MG nuclei are outlined by dotted-lines. **(b)** The thickness of the ONL in the healthy and RP patient retina was compared to the thickness of the INL in the same section, and the relative values are presented in the graph (data from 6 independent retinal sections from 2 eyes). **(c)** Prox1 intensity in Sox2-positive MG relative to other retinal neurons in the same image is presented in the graph (data from 12 independent retinal sections from 2 eyes). *** p<0.005; **** p<0.001 (Student’s t-test). **(d)** The distribution of Prox1 in the retinas of mice with heterozygous and homozygous *rd10* mutations in *Pde6b* gene (*Pde6b^rd10^*) was investigated by immunostaining. MG cells among Prox1-positive cells were determined by co-staining of Prox1 and Sox2. Boxed areas in top row are magnified in the next three rows. Sox2-positive MG nuclei are outlined by dotted-lines. **(e)** Prox1 intensity in Sox2-positive MG relative to other retinal neurons in the same image is presented in the graph (data from 3 independent litters). ** p<0.01; *** p<0.005; **** p<0.001 (Student’s t-test). (**f**) *Pde6b^rd10/rd10^*;*Glast-CreERT;R26^tdTom/+^*mice were intravitreally injected with AAV2-Ctrl Ab or AAV2-αProx1 at P10, followed by Tam injection to activate the CreERT, resulting in the expression of tdTom Cre reporter in MG. Mice were also injected with EdU at P34, P36, and P38 to label cells born during last 5 days before sample collection at P39. (**g**) The identities of tdTom-expressing MG cell lineage cells in mouse retinas were investigated by co-immunostaining for cell type-specific markers. Immunostaining images are provided in Extended Data Fig. 13e, except for the rPR marker, Rhodopsin. The birth of these cells between P34 and P39 was determined by EdU-labeling. Boxed areas in top row are magnified in the next rows. White arrowhead points EdU-labeled cell rPR nuclei and black arrowhead indicates EdU-labeled cell nucleus in the choroid. (**h**) Mean numbers of cells expressing corresponding markers in the retinal area are provide in the graph (data collected from 4 independent litters). (**i**) Visual acuities of the mice measured at the indicated postnatal days. Error bars in the graphs represent SEM (n=8; 5 independent litters).

Thus, we assessed the therapeutical potential of αProx1 in retinal degenerative diseases by infecting *Pde6b^rd^*^10^*^/rd^*^10^ mouse retina with AAV2-αProx1. We observed a significant decrease in Prox1 intensity in MG of *Pde6b^rd^*^10^*^/rd^*^10^ mouse retinas infected with AAV2-αProx1 compared to that in MG of AAV2-Ctrl Ab-infected *Pde6b^rd^*^10^*^/rd^*^10^ mouse retinas (Extended Data Fig. 13a,b).

Additionally, we noted a significant increase in the number of rPRs in the outer nuclear layer (ONL) in AAV2-αProx1-infected *Pde6b^rd10/rd10^*mouse retinas compared to those in uninfected or AAV2-Ctrl Ab-infected *Pde6b^rd10/rd10^*mouse retinas (Extended Data Fig. 13a,c). However, the numbers of TUNEL-positive apoptotic cells in AAV2-αProx1-infected *Pde6b^rd10/rd10^*mouse retinas were not significantly different from those in the others (Extended Data Fig. 13a,d), indicating that the increase of the rPRs was not resulted from the decrease of cell death.

Thus, we investigated the increase of the ONL thickness was related with the regeneration of rPRs by identifying EdU-labeled newborn cells in the mouse retinas. Significant numbers of EdU-labeled newborn Rhodopsin-positive rPRs were found in the ONL of AAV2-αProx1-infected *Pde6b^rd10/rd10^;Glast-CreERT;R26^tdTom/+^*mouse retinas (Fig. 4f,g). EdU incorporation was also observed in Sox2-positive MG or MGPCs and Pkcα-positive rod bipolar cells (rBCs) in those retinas, whereas the majority of EdU-labeled cells were microglia in AAV2-Ctrl Ab-infected samples (Fig. 4h; Extended Data Fig. 13e). Furthermore, those EdU-labeled rPRs and rBCs in AAV2-αProx1-infected *Pde6b^rd10/rd10^;Glast-CreERT;R26^tdTom/+^*mouse retinas express tdTom MG lineage cell marker (Fig. 4g,h; Extended Data Fig. 13e). These results suggest MG in AAV2- αProx1-infected *Pde6b^rd10/rd10^* mouse retinas not only expanded themselves but also differentiated into the retinal neurons.

Corresponding the gain of rPRs, the amplitudes of scotopic electroretinogram (ERG) a-waves were remarkably elevated in AAV2-αProx1-infected mice at postnatal day 35 (P35) in comparison to uninfected or *Pde6b^rd10/rd10^* littermate mice, suggesting that rPR activities were remained in the AAV2-αProx1-infected *rd10* mouse retinas (Extended Data Fig. 14a,b,c). Further, we observed that visual acuity of AAV2-αProx1-infected *Pde6b^rd10/rd10^* mice was significantly higher compared to that of uninfected or AAV2-Ctrl Ab-infected *Pde6b^rd10/rd10^*mice (Fig. 4i).

However, these effects were not persistent by P60, when the visual functions of AAV2-αProx1- infected *Pde6b^rd10/rd10^*mice were also impaired (Fig. 4i).

### Vision recovery in late onset retinitis pigmentosa mouse models by blocking Prox1 transfer

The transient vision recovery observed in AAV2-αProx1-infected *Pde6b^rd10/rd10^*mice was likely associated with the short lifespan of newly produced PRs, which also carry the *Pde6b^rd10^* mutation and degenerate in a week (Extended Data Fig. 15a,c). Therefore, the regeneration promoting effects of αProx1 might be more effective in retinas where degeneration takes for a longer period. We thus selected *Rp1^tvrm64/tvrm64^* (*Rp1^m/m^*) mice, which harbor homozygous missense mutations in the *retinitis pigmentosa 1* (*Rp1*) gene and start to lose rPRs from 2 month-old following to the shortening of outer segments and gradually lose their vision over 6 months ^40^ (Fig. 5a,b,g).

**Figure 5.**
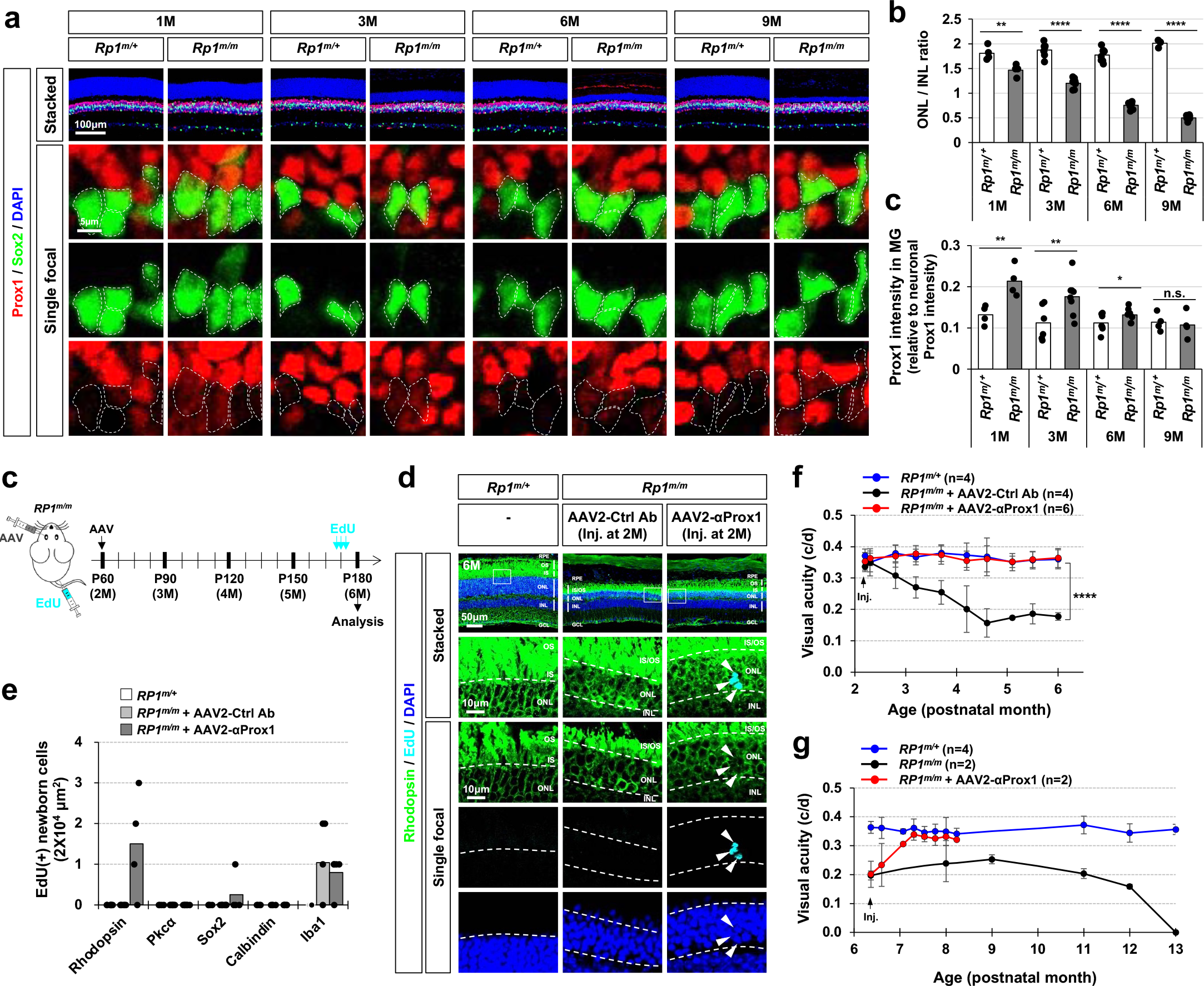
Vision recovery in late onset RP model mice by viral gene delivery of anti-Prox1 antibody. (a) The distribution of Prox1 in the retinas of mice with heterozygous and homozygous *tvrm64* mutation of *Rp1* gene (*Rp1^m^*) was investigated by immunostaining. MG cells among Prox1-positive cells were determined by co-staining of Prox1 and Sox2. Boxed areas in top row are magnified in the next three rows. Sox2-positive MG nuclei are outlined by dotted-lines. **(b)** Prox1 intensity in Sox2-positive MG relative to other retinal neurons in the same image is presented in the graph (data from 3 independent litters). ** p<0.01; *** p<0.005; **** p<0.001 (Student’s t-test). (**c**) *Rp1^m/m^* mice were intravitreally injected with AAV2-Ctrl Ab or AAV2-αProx1 at P60, and then with EdU at 9, 7, and 5 days before sample collection to label the cells born around the injection time points. (**d**) The identities of these EdU-labeled cells in mouse retinas were investigated by co-immunostaining for cell type-specific markers. Immunostaining images are provided in Extended Data Fig. 16e, except for the rPR marker Rhodopsin. Boxed areas in top row are magnified in the next rows. Arrowheads, EdU-labeled cells containing Rhodopsin. (**e**) Number of cells expressing corresponding markers (data collected from 2 independent litters). (**f**) Visual acuities of the mice infected with AAV at 2 postnatal months and measured at the indicated postnatal months. (**g**) Visual acuities of the mice infected with AAV at 6M and measured at the indicated postnatal months. Error bars in the graphs represent SEM. * p<0.05; ** p<0.01; *** p<0.005 (ANOVA).

Prox1 protein levels were observed to be higher inMG of *Rp1^m/m^* mice compared to their littermate *Rp1^m/+^*mice (Fig. 5a,c), consistent with findings in a retinitis pigmentosa (RP) patient and and *Pde6b^rd10/rd10^*mice (Fig. 4, a-e). The increased Prox1 expression in MG was mitigated by intravitreal injection of AAV2-αProx1 but not by AAV2-Ctrl Ab (Extended Data Fig. 16a,b). Notably, the thickness of the ONL was increased in *Rp1^m/m^* mice infected with AAV2-αProx1 compared to *Rp1^m/m^* mice infected with AAV2-Ctrl Ab (Extended Data Fig. 16a,c), despite no significant alteration in apoptotic cell numbers in the mouse retinas (Extended Data Fig. 16a,d). This increase in ONL thickness was associated, at least in part, with the regeneration of rPRs.

Significant numbers of EdU-labeled newborn cells in *Rp1^m/m^*mouse retinas infected with AAV2- αProx1 were Rhodopsin-positive rPRs, whereas EdU incorporation was not observed in littermate *Rp1^m/m^* mice infected with AAV2-Ctrl Ab (Fig. 5c-e; Extended Data Fig. 16e). The *Rp1^m/m^*mice infected with AAV2-αProx1 were able to maintain normal visual acuity at 6 months, whereas those treated with AAV2-Ctrl Ab experienced a gradual decline in vision (Fig. 5f). Furthermore, *Rp1^m/m^*mice infected with AAV2-αProx1 at 6 months showed visual acuity recovery close to that of *Rp1^m/m^* littermate mice at 8 months (Fig. 5g). Collectively, the blocking PROX1 transfer could be an effective therapeutic strategy for late onset retinal degenerative diseases, in which degeneration is relativel slower than the early onset types.

## Discussion

In the injured zebrafish retina, MG are reprogrammed to MGPCs, which reenter the cell cycle to renew themselves and then exit cell cycle to produce retinal neurons ^3^. In contrast, mouse MG barely turn into proliferative MGPCs after retinal injuries, although they are activated to exhibit different characteristics from resting MG ^6^. Therefore, the critical factors that induce MG reprogramming and proliferation may be missing from mammalian retinas. A strong candidate for this missing factor is Ascl1, which has been shown to be capable of triggering cell proliferation upon overexpression in the MG of injured mouse retinas ^10,11^. However, the mechanisms underlying the expression of *ascl1a* in fish MGPC and the suppression of *Ascl1* in mammalian MG remain unknown. In fly neurogenesis, the expression of fly Ascl1 homologs *scute* and *asense* is suppressed by the fly Prox1 homolog pros ^15^. This suggests that exogenous Prox1 accumulated in MG of injured mouse retina may be responsible for the suppression of *Ascl1* expression in the MG. However, *Ascl1* was not recovered in the MG of *Prox1^fg/fg^;Chx10-CreERT2* mouse retinas or AAV2-αProx1-infected mouse retinas, in which Prox1-depleted MG or MGPCs regained proliferative potential (Extended Data Fig. 12b; results for AAV2-αProx1-infected mouse retinas are not shown). The failure of *Ascl1* expression in those Prox1-depleted MGPCs is likely related to the upregaulation of Notch signaling, which was identified to repress the expression of Ascl1 ^41^.

Therefore, Prox1-depleted MG of injured mouse retina might be reprogramed into MGPCs independent of Ascl1 induction but through alternative mechanisms, including the Notch1-Hes1 pathway that is negatively regulated by Prox1 ^17^ (Fig. 3n).

Prox1 depletion could mediate other proneural transcription factors in MG for MGPC reprogramming in the absence of Ascl1 induction. Mouse Prox1 has been identified to suppress *Ccnd1* expression in NPCs of the cerebellar extra-ganglion layer ^35^. This resulted in the decrease of a proneural transcription factor, atonal bHLH transcription factor 1 (Atoh1), via the decrease of the Cdk4/Ccnd1 complex, which phosphorylates and stabilizes Atoh1 ^35^. Ccnd1 was found to be elevated in MGPC populations in the MNU-injured *Prox1^fg/fg^;Chx10-CreERT2* mouse retinas and AAV2-αProx1-infected *Glast-CreERT* mouse retinas (Fig. 3e,f,k,l,m; Extended Data Fig. 11d,e), suggesting an increase of Atoh1 in these cells. However, Atoh1 was not detectable in those Prox1-depleted mouse retinas (data not shown). Hence, the reprogramming of MG into MGPC by depletion of Prox1 in those mouse retinas might involve other proneural transcription factors, including Neurod1, which was found in cluster #20 at a significant number (Extended Data Fig. 12b). However, future studies need to address the specific proneural transcription factors that play a critical role in the transition of MG to MGPCs.

Prox1 has been identified as a critical player in the development of HCs in the mouse retina ^19^. However, we could not find the transdifferentiation of Prox1-ennriched MG to HCs or other retinal neurons in injured or disease mouse retinas (Fig. 3k and 4h). We also found no evidence of the conversion of MG to HCs upon ectopic expression of *Prox1* in the MG of *Glast-CreERT;R26^tdTom/+^* mouse retinas (data not shown), implicating that Prox1 in mouse MG cannot induce spontaneous transdifferentiation to retinal neurons regardless of its sources. Given that epigenetic modifications are necessary for retinal regeneration from Ascl1-overexpressing MG ^10^, these results suggest that changing the epigenetic context of MG to one that is more similar to developing RPCs would be necessary for Prox1-mediated conversion of MG to HC in the adult mouse retina.

Our findings indicate that deleting *Prox1* specifically in BCs was adequate to diminish Prox1 levels in MG, thereby enabling their reprogramming into MGPCs, despite other retinal neurons such as HCs and A2 ACs still expressing Prox1 (Fig. 2, a – e; Extended Data Fig. 5a,b). Additionally, sequestering extracellular Prox1 using αProx1 could reduce Prox1 levels in MG, leading to their conversion into MGPCs in the injured mouse retina without affecting endogenous *Prox1* expression in MG (Fig. 2f,g). These observations imply the existence of a threshold Prox1 level necessary to impede injury-induced MGPC reprogramming of MG. Therefore, Prox1 levels in MG may not reach this threshold in these mouse retinas (Fig. 2) and the injured zebrafish retina (Extended Data Fig. 7), enabling the conversion to MGPC from MG without the interference by the basal Prox1. However, the precise threshold level remains unknown at present.

Not only do MG extend processes to inner and outer retinal surfaces to form barriers, they also branch into the outer and inner plexiform layers (OPL and IPL) to modulate the synaptic transmission of retinal neurons ^42^. Thus, MG may capture Prox1 secreted from BCs into the OPL and IPL, where BCs form pre- and post-synaptic connections, respectively (Fig. 2a, model diagram). However, we found no changes in the Prox1 levels in ACs or HCs, which form synapses with BCs in the OPL and IPL, respectively, in the retinas of *Prox1^fg/fg^;Chx10-CreERT2* mice (Extended Data Fig. 5a,b), while we did find a significant decrease in Prox1 levels in the MG in these retinas (Fig. 2b,d). Therefore, extracellular Prox1 moves preferentially into MG compared to ACs and HCs in the mouse retinas. Orthodenticle homeobox 2 (Otx2) has been found to move short distances across synapses between PRs and BCs ^43^ and long distance from the choroid plexus to cortical parvalbumin (PV) interneurons ^44^. In the latter case, the specific binding of Otx2 to chondroitin-4-sulfate, which is strongly enriched in the perineural net surrounding PV interneurons, increases the selectivity of Otx2 transfer ^45,46^. A similar process may also be at work in MG, which may trap Prox1 by expressing specific types of heparan or chondroitin sulfate proteoglycans that are lacking in other retinal neurons. Interestingly, enzymes involved in introducing chondroitin sulfate (*Chst1* and *Chst5*) and heparan sulfate (*Ext1*, *Ext2*, *Extl1*, *Extl2*, *Extl3*, *Ndst1*, *Ndst2*, *Ndst3*, and *Ndst4*) are commonly upregulated between 6 and 12 hours after NMDA injury in the mouse MG lineage ^6^. However, levels of these homologous genes are not changed in the injured zebrafish retina ^6^, where Prox1 levels remain unchanged in MG after injury (Fig. 1b,c). Given the selective, but redundant, roles of these enzymes, understanding the selectivity of the transfer of Prox1 into MG will require future studies to identify which enzymes actually supply these specific sugar codes to MG.

The ONL thickness and scotopic ERG responses of *Rp1^m/m^*mice infected with AAV2- αProx1 were not completely restored to normal levels (Extended Data Fig. 17a,b,c), despite achieving complete recovery in visual acuity (Fig. 5f). These results suggest that the efficiency of retinal regeneration induced by AAV2-αProx1 infection was insufficient to fully restore the rPRs in *Rp1^m/m^*mice. Instead, the mice exhibited an increase in cPR numbers and photopic ERG responses, which are attributed to the function of cPRs (Extended Data Fig. 17d,e,f; Extended Data Fig. 18). In patient with RP, the degeneration of cPRs follows the degeneration of rPRs before they lose vision completely, as the survival of cPRs depends on factors produced by rPRs. One such pathway involves the expression of rod-derived cone viability factor (RdCVF) in rPRs, which supports the survival of cPRs ^47^. Therefore, the partial recovery of Rhodopsin-expressing rPRs in *Rp1^m/m^* mice infected with AAV2-αProx1 may be sufficient to provide the protective factors to cPRs above a critical threshold concentration, thereby promoting their survival.

## Methods

### Human samples

Human donor eyes (bank tissue ID are 22-1465 [RP patient, 79 years-old] and 22-1587 [healthy donor, 83 years-old]) were collected in collaboration with the Lions Gift of Sight Eye Bank according to a standardized protocol of the bank. Briefly, eyes were procured within a 12-hour post-mortem interval. To determine the ocular phenotype relative to disease and healthy aging, ophthalmoscopic analysis was performed by a team of retinal specialists and ophthalmologists at the Lions Gift of Sight Research Team. This diagnosis was then compared to medical records.

The isolated eyes were fixed in phosphate buffered saline (PBS) containing 4% paraformaldehyde (PFA) and delivered to the laboratory in a container filled with icepacks within 36 hours. The fixed eyes were dissected to quadrants for subsequent cryopreservation in OCT (Tissue-Tak). The frozen eye sections were used for the immunohistochemical analyses described in the following. All procedures were done according to the protocol approved by KAIST Institutional Review Board (IRB-22-551).

### Mice

*Prox1^tm^*^1^*^.1Fuma^* (*Prox1^fg^*)^18^, *Dkk3^tm1Tfur^* (*Dkk3-Cre*)^48^, *Tg(Chx10-cre/ERT2)G7Tfur* (*Chx10-CreERT2*), *Tg(Slc1a3-cre/ERT)1Nat/J* (*Glast-CreERT*)^26^, *Tg(Crx-cre/ERT2)1Tfur* (*Crx-CreERT2*)^24^, *B6.CXB1- Pde6b^rd^*^10^*/J* (*Pde6b^rd^*^10^), and *C57BL/6J-Rp1^tvrm^*^64^*/PjnMmjax* (*Rp1^m^*)^40^ mice were maintained and bred in a specific pathogen-free mouse facility of KAIST Laboratory Animal Resource Center. For intravitreal injections, mice were anesthetized with isoflurane (1.5% induction and 1.0% maintenance). Samples loaded into a Hamilton syringe equipped with a blunt 33-gauge needle were then injected into the intravitreal space of the mouse eye. All mouse experiments were performed according to the protocol approved by KAIST institutional animal care and use committee (KAIST IACUC 13-130 and KA2019-14).

### Zebrafish

AB (wild-type) and *Tg(gfap:EGFP)* zebrafish (*Danio rerio*) strains were maintained at 28.5±1°C with a 14/10-hour light/dark cycle. To injure the fish retina using MNU, adult zebrafish (6-12 months of age) were incubated in fresh media containing MNU (150 mg/l) and 10 mM sodium phosphate (pH 6.3) for 1 hour and then washed with system water ^49^. The MNU-treated zebrafish were maintained under standard husbandry conditions. To label proliferating cells, the zebrafish were anesthetized with 0.04% Tricaine (ethyl 3-aminobenzoate methanesulfonate salt, #A5040, Sigma, USA) in system water and given *i.p.* injections of 20 µl of 50 mg/ml EdU (dissolved in PBS) at the specified times. All zebrafish care and husbandry were performed in compliance with the guidelines of the Korea Research Institute of Bioscience and Biotechnology (KRIBB) IACUC (KRIBB-AEC-20235).

### Immunohistochemistry

Proteins in the retinal sections and culture slides were detected by immunostaining, respectively, as described previously ^43^. Sections (20 µm) of frozen zebrafish and mouse eyes were incubated in a blocking solution (PBS including 1% normal donkey serum and 0.1% Triton X-100) at room temperature for 2 h. The sections were further incubated in blocking solution (without Triton X- 100) containing primary antibodies at 4°C for 16 h, and then with fluorophore-conjugated secondary antibodies that recognize the primary antibodies. The information on antibodies used in this study is provided in the Table 1. Fluorescent images were then obtained using Olympus FV1000 and FV3000 confocal microscopes.

### Quantification of immunostaining signals

Single focal confocal microscopic images were converted to grayscale and utilized for quantifying fluorescent signal intensity. Subsequently, the cell images were adapted to facilitate nucleus segregation analysis. Prox1 signals in MG were captured within Sox2+ areas. For acquiring Prox1 signals in BCs and ACs, the outermost 2-3 layers of the inner nuclear layer (INL) were delineated as the BC region, while the innermost single layer of the INL was designated as the AC region.

These regions were delineated using the lasso tool in Photoshop and then exported. GFP images were binarized using the Otsu algorithm in Python, and the nuclei on DAPI images containing

>50% of binary GFP in nucleus areas were selected for quantifying fluorescence intensity.

Prox1 images and the modified target cell images were imported into Python. Nucleus segregation of both Prox1 and target cell images was conducted using the StarDist algorithm ^50,51^. The Prox1 image was normalized by the mean value of Prox1 fluorescent intensity in segregated Prox1+ nuclei. Segregated target nuclei with >200 pixel counts were utilized to quantify Prox1 fluorescent intensity. To minimize spill-over Prox1 fluorescence from neighboring Prox1+ cells, overlapping Prox1+ nuclei with the selected single target nucleus were identified. If the overlapping Prox1+ nucleus was not a subset of the target nucleus, the Sørensen–Dice coefficient (SDC) between the target nucleus and each overlapping Prox1+ nucleus was calculated to define overlapping regions with an SDC < 0.5 as ’spill-over’ regions. The target nucleus region was then adjusted by excluding these spill-over regions

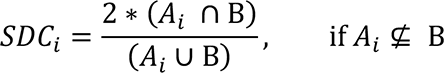

Where A is i-th neighbor Prox1^+^ nucleus region and B is a target nucleus region.

The pixel intensity matrix of the corrected target nucleus region was then extracted from normalized Prox1 images. The median value of the pixel intensity matrix was computed and represented as the Prox1 intensity value of a single target nucleus. After obtaining median values from all target nuclei, the mean of these median values was considered as the Prox1 intensity value of one single-focal image, which was used for subsequent analysis and plotting.

To quantify GFP signal intensity, segregated target nuclei with >200 pixel counts were utilized. Histogram equalization was applied to each GFP image using the ’cv2.equalizeHist()’ function for comparison with *Prox1^fg/+^;Dkk3-Cre* mouse retinal images. GFP intensity was measured using the same method as for Prox1 intensity measurement, with the exception of spill- over correction

### RNAscope in situ hybridization

RNAscope *in situ* hybridization was conducted to detect mRNA expression in mouse retinal sections following the manufacturer’s instructions. For co-staining of Prox1 mRNA and Sox9 protein, the RNA-Protein Co-Detection Ancillary Kit (Cat. #323180) and a Sox9 antibody were employed. The Multiplex Fluorescent V2 Assay utilized to probe Prox1 (Probe- Mm-Prox1-C1, Cat. # 488591, ACDbio), while Sox2 protein was detected by immunostaining. Subsequently, the fluorescence signals from the Prox1 probe and Cy3-labeled donkey anti-rabbit secondary antibody were visualized using confocal microscopy

### Quantification of Prox1 RNAscope signals

Single-focal target cell images and RNAscope images were imported on Python and nucleus segregation was performed using Stardist software. *Prox1* RNAscope signals inside segregated target nuclei (>200 pixel-counts) were exported as gray-scaled images for quantification. Imported *Prox1* RNAscope images on ImageJ were converted to binary image by using ‘Threshold’ tool (minimum threshold value = 1, maximum threshold value = 255, and activated- ‘dark background’ option). The particle size and number of binary *Prox1* RNAscope signals were measured through ‘Analyze Particles’ tool with activated- ‘Include holes’ option. The particle size and number were divided by the number of segregated nuclei.

### Recombinant Prox1 protein production and intravitreal injection

The PROX1 cDNA was subcloned into pFASTBAC1 to express PROX1 with an N-terminal FLAG-tag in Sf9 cells by the baculovirus expression system (Invitrogen). The PROX1-FLAG protein was then purified on M2 agarose (Sigma), which captures FLAG-tagged proteins as described previously ^52^. The purified proteins were diluted in sterilized PBS and 1 µl of the protein/PBS solution loaded into a Hamilton syringe equipped with a blunt 33-gauge needle for injection into the intravitreal space of the mouse and zebrafish eyes. Contralateral eyes received a sham injection of PBS alone and served as controls.

### Isolation of αProx1 clone

An antibody that has an affinity for Prox1 was screened from a phage library of chicken immunoglobulin genes by YntoAb Inc. (Korea). A clone (1A11) that exhibited strong affinity to the antigen was chosen for further analyses *in vitro* and *in vivo*. The immunoglobulin gene clone was prepared in a single-chain fragment (scFv) and fused with hemagglutinin plus 6 histidine sequences (6XHis-HA) to allow it to be detected by the 6XHis-HA tag. The clone was also fused with the signal sequence of human interleukin-2 (IL2) before introduction into the genome of adeno-associated virus (AAV) for the expression of the scFv antibody in mouse retina.

### Single cell RNA analysis: cDNA library synthesis

Freshly prepared samples were resuspended in resuspended in the resuspension buffer (PBS containing 0.04% BSA and 0.5U/ml RNase inhibitor). Dissociated cells were then filtered through a 50-ml cell filter. Cells (∼10,000) were loaded into a 10x Genomics Chromium Next GEM Single Cell system (10x Genomics, CA). Single cell cDNA libraries were prepared using the Chromium Next GEM Single Cell 3ʹ Kit according to the manufacturer’s instructions. Synthesized cDNA libraries were pooled and sequenced on Illumina pair ended sequencing system.

### Single cell RNA analysis: quality control and visualization of scRNA-seq data

Raw sequencing files were converted to FASTQ files by the cellrnager mkfastq software, and then to gene-cell matrices by the cellranger count command. Gene-cell matrices were analyzed by the Seurat package to visualize scRNA-seq data ^53^. Cells that have >20% mitochondrial counts or less than 1,000 nFeature value were excluded, and the filtered cells was processed through Seurat integration pipeline to cluster them on UMAP using t-distributed stochastic neighbor embedding (tSNE) ^54^. The clusters visualized on UAMP were annotated by cell type-specific markers. Retinal cells expressing specific genes were quantified on RNA slot. Mean cross bar for vlnplot was processed by stat_summary and mean crossbar data was merged with vlnplot data on Adobe Photoshop 2021.

### Single cell RNA analysis: RNA velocity analysis

Spliced/unspliced dataset was calculated by velocyto package to obtain loom files. The loom files were then loaded on R studio through the velocyto.R and SeuratWrappers R libraries to convert into Seurat class. Spliced slot was normalized by SCTransfrom to pull the count data and UMAP reduction was proceeded. Velocity value was calculated by Runvelocity and was projected onto embeddings coordinates of integration-seurat-object.

### SMART-seq: Single cell cDNA synthesis and sequencing library generation

Single-cell RNA sequencing (scRNA-seq) libraries were prepared using the Smart-seq2 protocol with few modifications ^55^. Single-cells were sorted using a BD FACSAria Fusion (BD Biosciences) into 96 well PCR plates (Thermo Scientific) containing 2μl of lysis buffer (0.1% Triton X-100, 1U/μl RNase Inhibitor (Enzynomics), 0.25μM oligo-dT30VN primer) and stored at -80℃. The plates were thawed on ice. Revers transcription (20U/µl Maxima H minus transcriptase, 1M Betaine, 5mM MgCl2, 1μM template switching oligo, additional 0.8U/μl RNase Inhibitor), template switching reaction and PCR pre-amplification (KAPA HiFi HotStart (Roche), 18 cycles) were performed according to the protocol. The PCR products in each well was cleaned up using 0.6× SPRI beads (2% Sera-Mag Speed Beads (Cytiva), 1M NaCl, 10mM Tris-HCl pH 8.0, 1mM EDTA, 0.01% NP40, 0.05% Sodium Azide, 22% w/v PEG 8000). The quality of cDNA libraries was assessed by quantitative PCR with a primer pair of GAPDH—a housekeeping gene (forward: 5′- GTCGTGGAGTCTACTGGTGTCTTCAC-3′; reverse: 5′- GTTGTCATATTTCTCGTGGTTCACACCC-3′). 50–100 pg of each cDNA library were used to generate the Illumina sequencing library using EZ-Tera XT DNA library preparation kits (Enzynomics). After the final PCR amplification, samples were pooled and cleaned by MinElute PCR purification kit (Qiagen). Using 0.3× and 0.6× SPRI beads, fragments of ∼200bp length were selected as library and confirmed by High Sensitivity DNA ScreenTape Analysis (Agilent). Pooled and size-selected sequencing libraries were sequenced on an Illumina NextSeq 550 instrument using a High Output kit (Illumina) with 38-bp paired-end reads setting.

### SMART-seq: scRNA-Seq data analysis

Sequencing reads from single-cell RNA-Seq libraries were aligned to the mouse reference genome (version mm10 from the UCSC) using RSEM (version 1.3.1) with STAR (version 2.7.9a) aligner with default parameters for paired-end reads ^56^. The matrix of raw counts was transformed into Seurat object (version 4.0.2) for the downstream analysis. SMART-seq data was then analyzed on Seurat ^57,58^ and monocle3 ^59,60^. Each raw count seurat objects were normalized by SCTransform and were integrated. Integrated Seurat object was converted to cell data set object compatible with monocle3 by as.cell_data_set(). Converted integrated-object was performed by common workflow (pre-processing, dimensionality reduction and clustering cells). To analyze trajectory analysis, the cell farthest from AAV2-αProx1+MNU sample on UMAP was specified as the root node by order_cells(). Gene expression figure by pseudotime was plotted by plot_cells() command. After cluster number in monocle object was inserted into integrated Seurat object, heatmap was plotted by DoHeatmap() command.

### Visual acuity test by OptoMotry

Mouse visual acuity was measured by the OptoMotry system (Cerebral Mechanics) as described previously ^61^. Mice were adapted to ambient light for 30 min and placed on a platform 60 cm above the floor and surrounded by computer monitors displaying black and white vertical stripe patterns. The mice stopping movement to begin tracking the stripe movements with head-turn was counted as a successful visual detection. The detection thresholds were then obtained from the OptoMotry software.

### Electroretinogram (ERG)

Mice were kept in the dark for 12 h before scotopic ERG recording and anesthetized with 2,2,2- tribromoethanol (Sigma). A gold-plated objective lens was placed on the cornea and silver-embedded needle electrodes placed at the forehead and tail after the pupils were dilated by 0.5% tropicamide. The ERG recordings were performed using a Micron IV retinal imaging microscope (Phoenix Research Labs) and analyzed by Labscribe ERG software according to the manufacturer’s instructions. A digital bandpass filter ranging from 0.3 to 1,000 Hz and stimulus ranging from −2.2 to 2.2 log(cd·s m^−2^) were used to obtain scotopic ERG waves, and a filter ranging from 2 to 200 Hz and stimulus ranging from 0.4 to 2.2 log(cd·s m^−2^) with 1.3 log(cd·s m^−2^) background were used to obtain photopic ERG waves.

### Statistical analysis

Statistical tests were performed in Prism Software (GraphPad; v5.0). All data from statistical analyses are presented as the mean ± standard error (STE). Comparisons between two groups were made by unpaired Student’s t-test, and the differences among multiple groups were determined by analysis of variance (ANOVA) with Tukey’s post-test. P-values < 0.05 were considered significant.

## Acknowledgements

We are grateful to Drs. Takahisa Furukawa and Jeremy Nathans for generously providing Dkk3- Cre, Chx10-CreERT2, Crx-CreERT2 and Glast-CreERT mice, which were instrumental for our research. We also extend our sincere appreciation to those who donated their eyes for this study, enabling us to further our understanding in this field.

## Author contributions

E.J.L., M.K., and J.W.K. wrote the manuscript; E.J.L., M.K., S. P., J.H.S., H.J.C., J.A.P., K.P., D.L., H.J., and J.H.K. designed and performed the experiments and analyzed the data; F.M. provided an experimental model; S.Y.K., J.K., H.Y., J.S.L., and J.W.K. supervised the project.

## Data and materials availability

The scRNA-seq data are available at KoNA at Korea Bioinformation Center (https://www.kobic.re.kr/kona/; Accession ID PRJKA220468). All other data are available in the main manuscript or the supplementary materials. Materials requests should be directed to J.W.K..

## Supplementary Information

**Extended Data Fig. 1.**
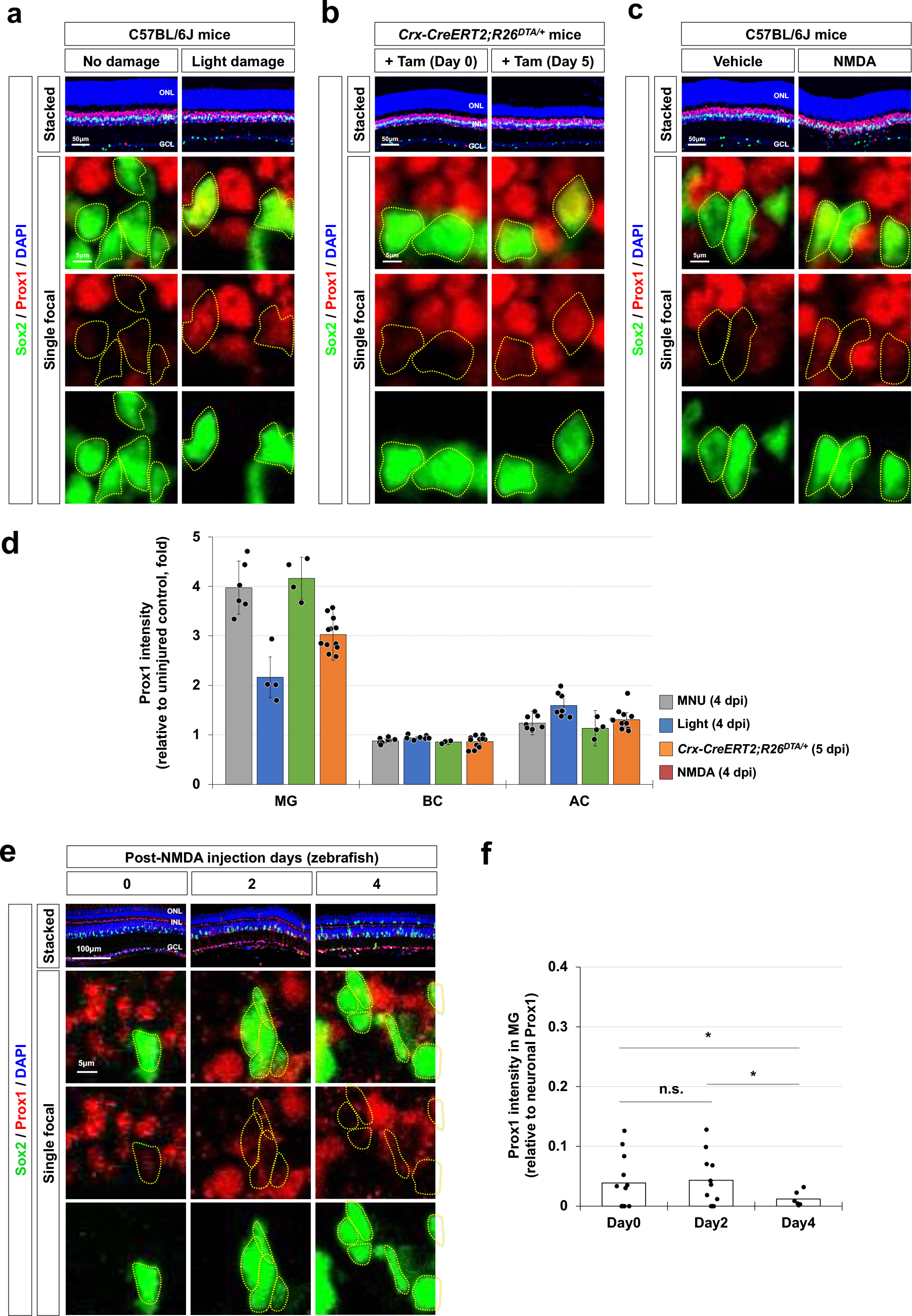
Upregulation of Prox1 in the MG of injured mouse retinas. (a) P30 C57BL/6J mice, dark-adapted and with dilated pupils, were exposed to light (100,000 Lux) for 3 hours. (**b**) P30 *R26^DTA/+^;Crx-CreERT2* mice were administered Tam (75 mg/kg) to activate CreER^T2^, facilitating the deletion of the LSL cassette to express diphtheria toxin A (DTA) in the photoreceptors. **(c)** P30 C57BL/6J mice were intravitreally injected with 0.5 µl of PBS (vehicle) or PBS containing NMDA (100 mM). After 7 (a) and 4 (b and c) days, the eyes were isolated for immunostaining to assess Prox1 expression. Sox2-positive MG cell nuclei are outlined by dotted lines. **(d)** Prox1 intensities in MG and ACs were normalized to that of BCs within the same images and their relative intensities are presented in the graph (data from 3 independent litters). **(e)** Zebrafish retinas, aged 6 months, were injured via intraocular injection of 2 µl NMDA (100 mM). Distribution of Prox1 in these fish retinas was then assessed by immunostaining. **(f)** The graph illustrates Prox1 intensity in MG relative to that in other retinal neurons within the same image (data from 3 independent litters). * p<0.05; n.s., not significant (p > 0.05).

**Extended Data Fig. 2.**
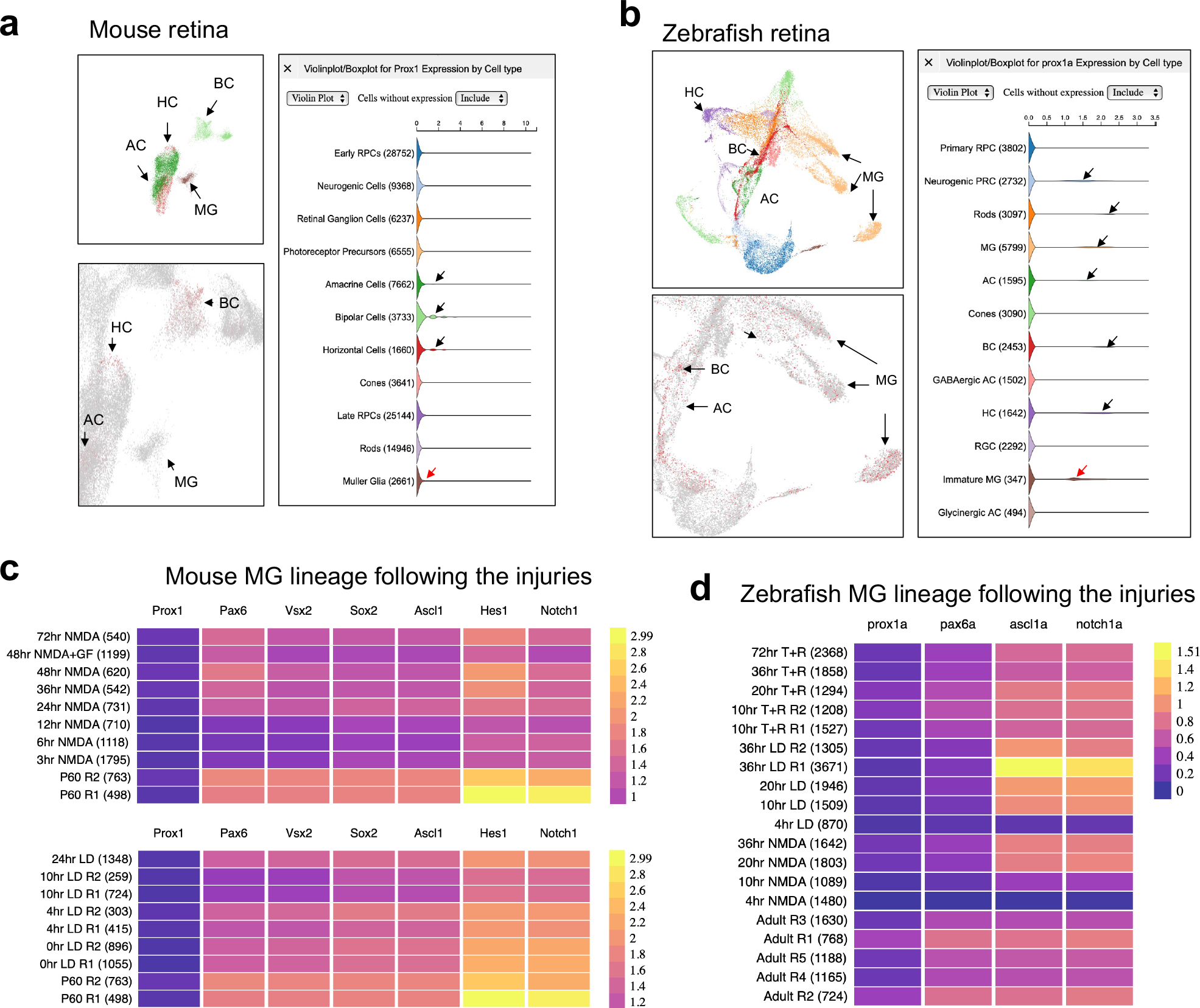
**Insignificant changes in Prox1 expression in MG of injured mouse and zebrafish retinas**. The presence of Prox1-expressing cells in mouse and zebrafish retinas was determined by analyzing single-cell RNA sequencing (scRNA-seq) data available at https://proteinpaint.stjude.org/F/2019.retina.scRNA.html. **(a)** In mouse retina, *Prox1* mRNA was detectable in BC, HC, and AC at significant level (black arrowheads); however, it was barely detectable in MG (red arrowheads). **(b)** In zebrafish retina, *prox1a* mRNA was broadly detectable in neurogenic RPC, rods, MG, AC, BC, HC and immature MG. **(c)** Analysis of purified MG lineage cells from mouse retinas subjected to NMDA damage or intense light damage (LD) revealed a scarce level of *Prox1* mRNA, which did not show significant changes following the injuries. **(d)** Similarly, in purified MG lineage cells from zebrafish retinas exposed to NMDA or LD, *prox1a* mRNA exhibited a slight decrease after the injuries.

**Extended Data Fig. 3.**
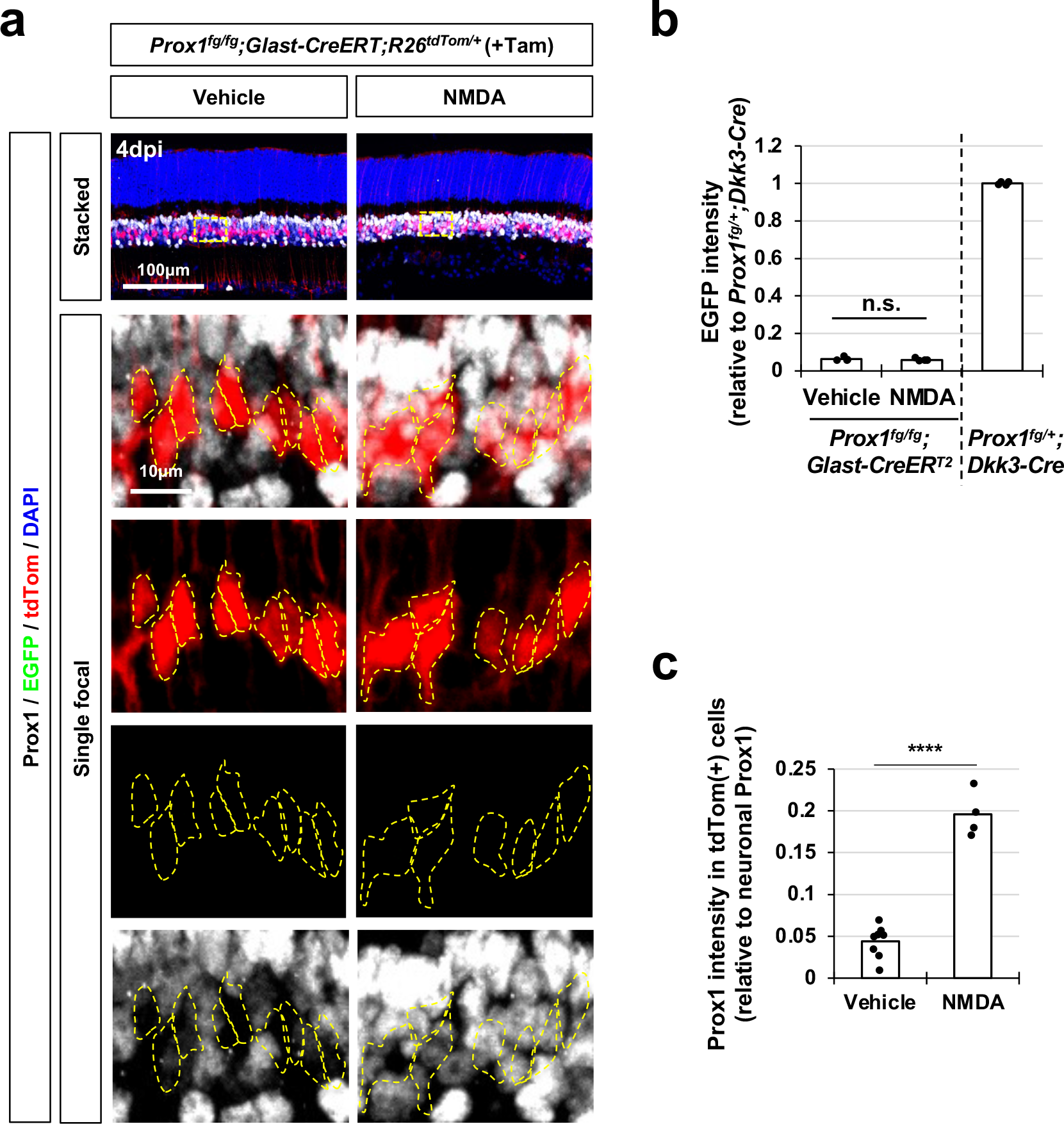
NMDA-induced accumulation of Prox1 protein in MG of *Prox1-cko* mice. (a) *Prox1^fg/fg^;Glast-CreERT;R26^+/tdTom^*(*Prox1-cko*) littermate mice were administered Tam at P21 and P22 to activate CreERT recombinase, leading to the deletion of Prox1 in the MG population prior to NMDA injection at P26. The endogenous expression of Prox1 in MG was indirectly determined by assessing EGFP expression in the *Prox1* gene locus. Boxed areas in top row are magnified in the next rows. The tdTom-positive cell bodies are outlined by dotted lines. **(b)** EGFP intensity of tdTom-positive cell body area was measured and the relative intensity to EGFP intensity of *Prox1^fg/fg^;Dkk3-Cre* mouse retina was presented in the graph (data from 2 independent litters). **(c)** The graph illustrates Prox1 intensity in MG relative to other retinal neurons within the same image. n.s., not significant; **** p<0.001 (Student t-test).

**Extended Data Fig. 4.**
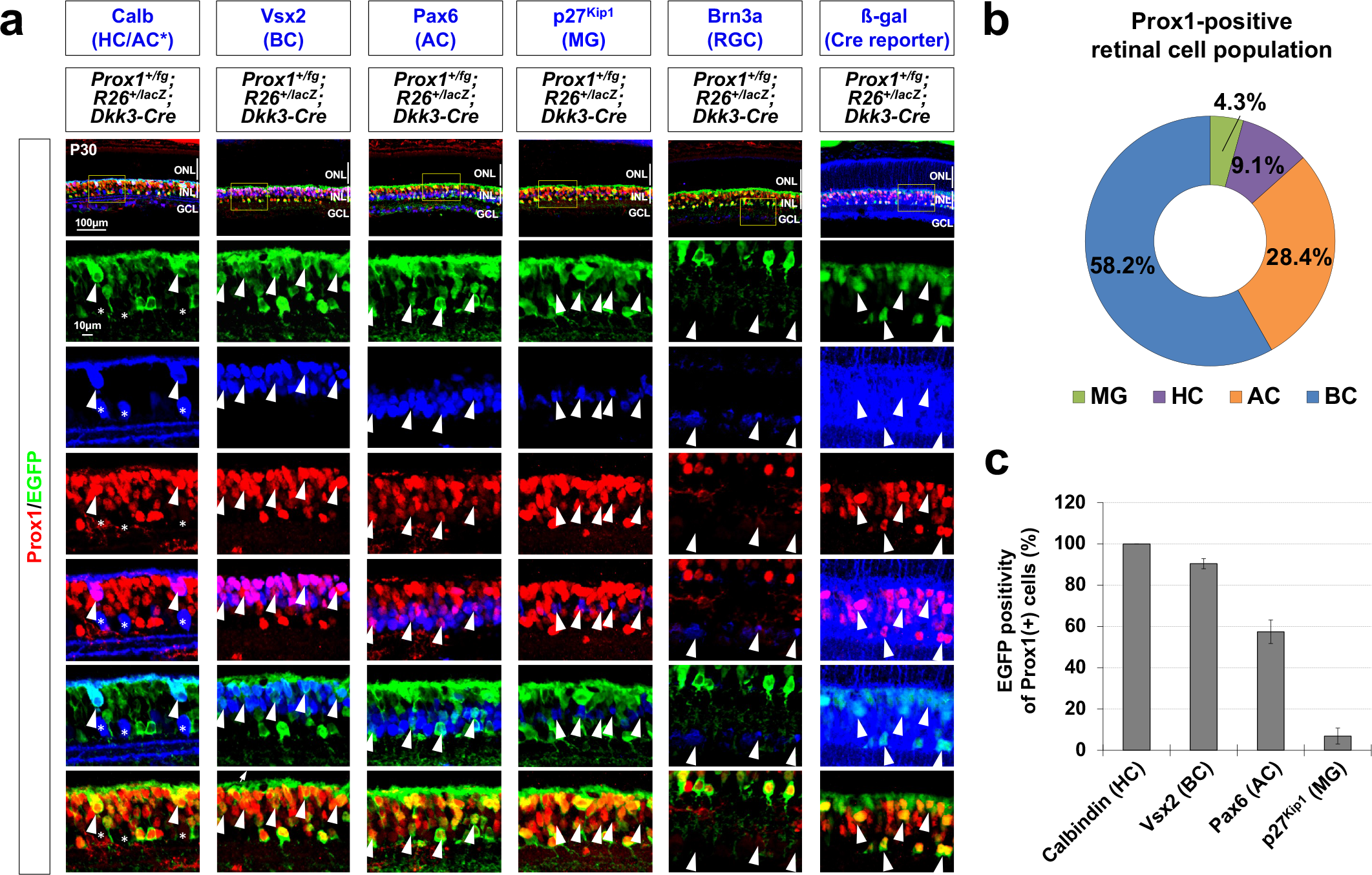
*Prox1* gene expression in HC, BC, and AC population in mouse retina. (a) The distribution of Prox1 and EGFP in the P30 *Prox1^+/fg^;Dkk3-Cre* mouse retina was investigated. The identification of Prox1- and EGFP-expressing cells in the retinas was conducted through co-immunostaining with corresponding markers for each retinal cell type. Cells undergoing Cre-dependent DNA recombination were visualized by immunostaining for ß- galactosidase (ß-gal) expressed in the *R26^lacZ^* locus. Boxed areas in the images in the top row are magnified in the bottom rows. Arrowheads point Prox1(+);Marker(+) cells. (**b**) Numbers of cells expressing corresponding markers in the retinas of P30 *Prox1^+/fg^;R26^+/lacZ^;Dkk3-Cre* mice counted and their ratio to total Prox1-expressing cell number was presented in the graph. **(c)** EGFP positivity of each Prox1-expressing retinal cell subset. Error bars represent SEM (n=6, 3 independent litters).

**Extended Data Fig. 5.**
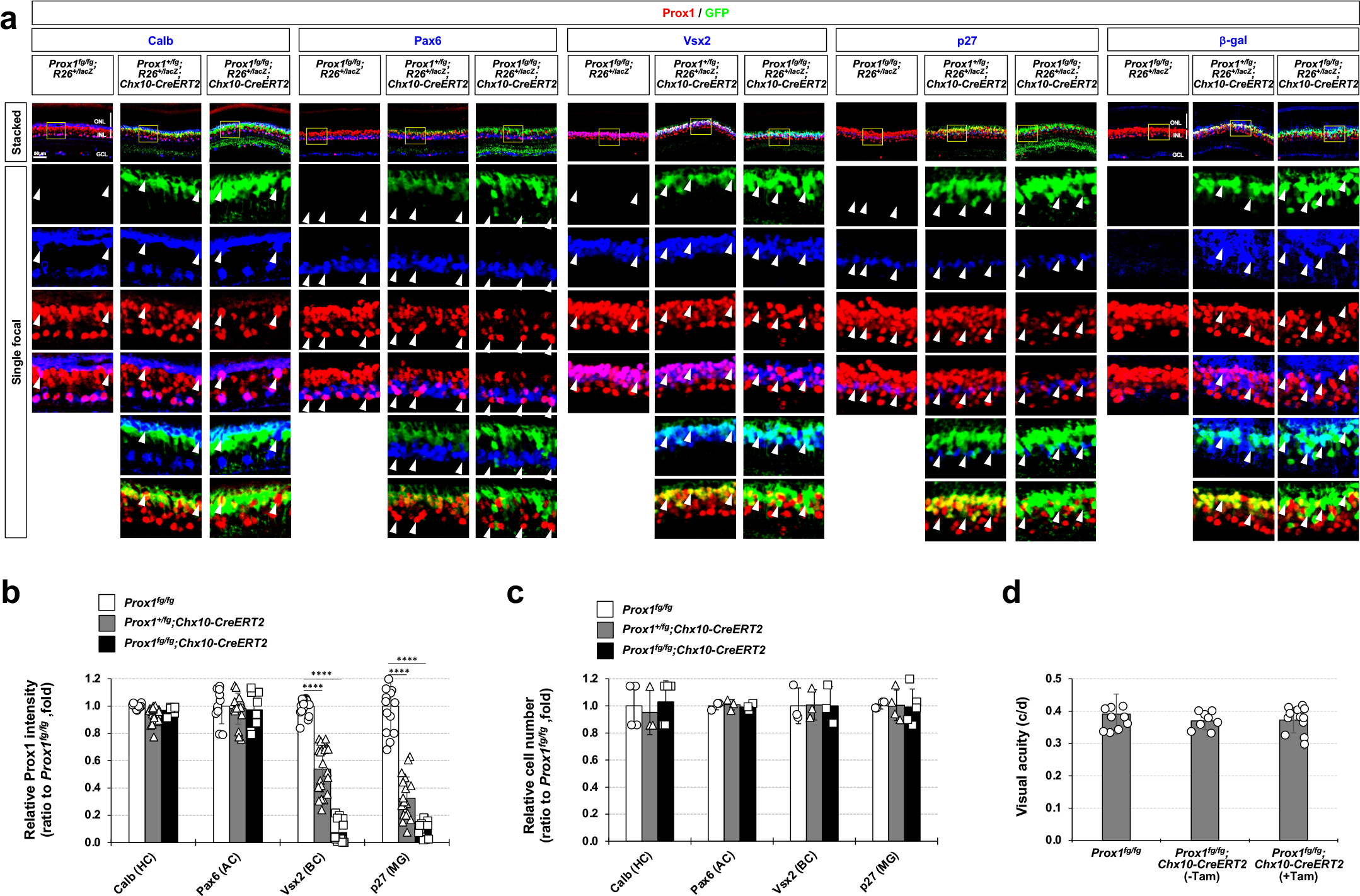
Deletion of *Prox1* in the BC population of *Prox1^fg/fg^;Chx10-CreERT2* mouse retinas. (a) Mice with the indicated genotypes were injected with Tam at P21, P22, and P23 to activate Chx10-CreERT2 recombinase, which is selectively expressed in BC population. The CreERT2-dependent expression of EGFP in the *Prox1^fg^* knock-in allele was assessed by confocal microscopy of eye sections from P40 mice. Cells underwent Cre-dependent DNA recombination were visualized by immunostaining for ß-gal Cre reporter. Boxed areas in the images in the top row are magnified in the bottom rows. Arrowheads point Prox1;Marker(+) cells. **(b)** The graph illustrates Prox1 intensities in HC, AC, BC, and MG relative to those in *Prox1^fg^* mice (data from 4 independent litters). * p<0.05; ** p<0.01; *** p<0.005 (ANOVA). **(c)** Number of cells expressing corresponding markers in the mouse retinas were counted and their ratio to the number in the retinas of P30 *Prox1^fg/fg^* littermates are presented (data from 4 independent litters). **(d)** Visual acuities of the mice were measured as described in Methods (data from 4 independent litters).

**Extended Data Fig. 6.**
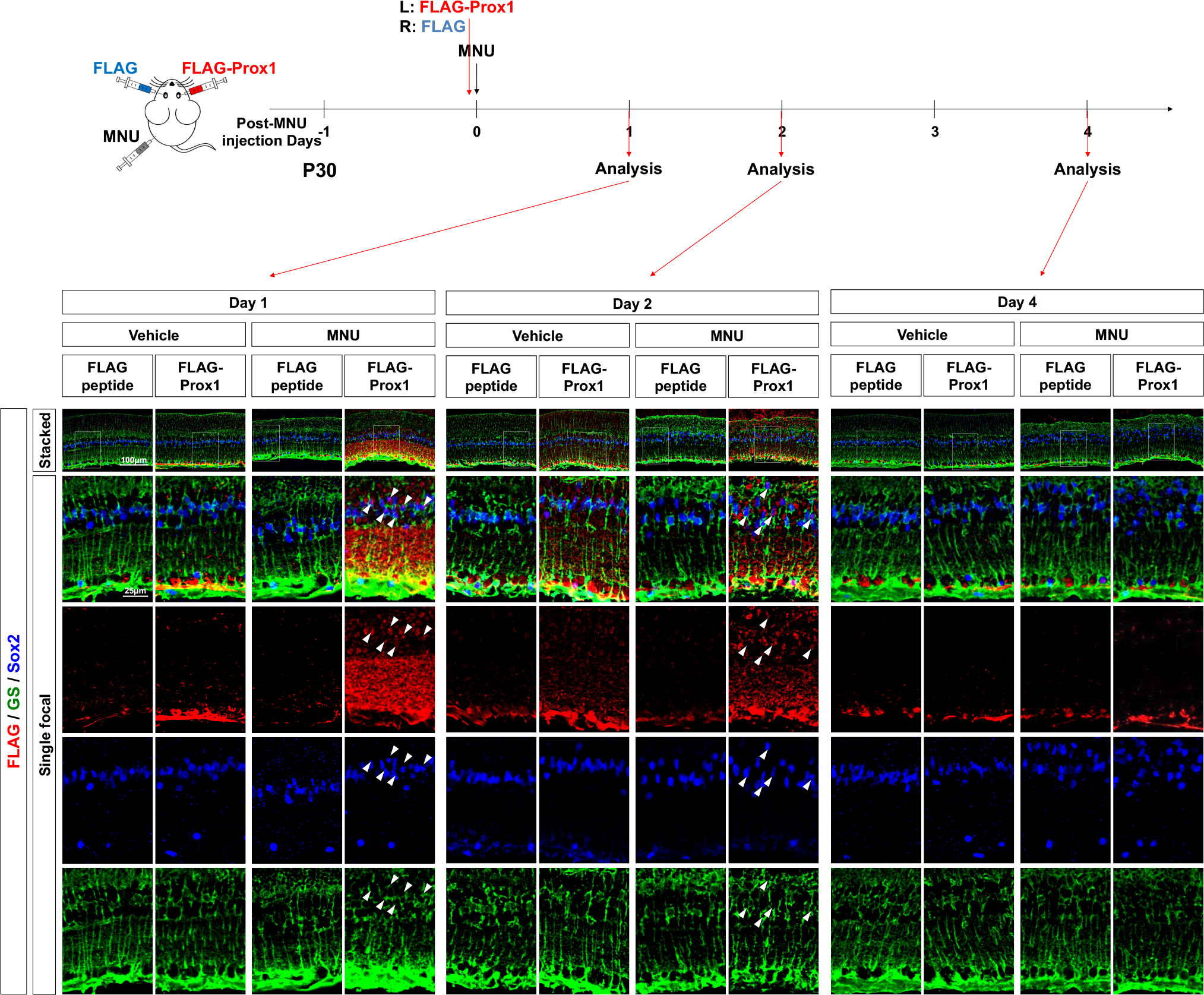
Delivery of Prox1 protein to the injured mouse retina. Equal volumes (1 µl) of FLAG peptide (10 ng/µl in PBS; administered to left eyes) and recombinant FLAG-Prox1 protein (20 ng/µl in PBS; administered to right eyes) were injected into the intravitreal spaces of P30 mouse eyes. This injection occurred 1 hour after intraperitoneal administration of vehicle (PBS containing 0.05% acetic acid) or MNU (60 mg/kg). The mouse eyes were collected at 1 day (Day 1; P31), 2 days (Day 2; P32), or 4 days (Day 4; P34) post-protein injection, and the presence of the injected FLAG-Prox1 proteins in the mouse retinas was evaluated by immunostaining using an anti-FLAG antibody (red). Penetration of the proteins into MG cells was identified through co-immunostaining with the anti-FLAG antibody and antibodies against MG cell markers, GS and Sox2. Arrowheads point FLAG-positive;Sox2-positive MG nuclei.

**Extended Data Fig. 7.**
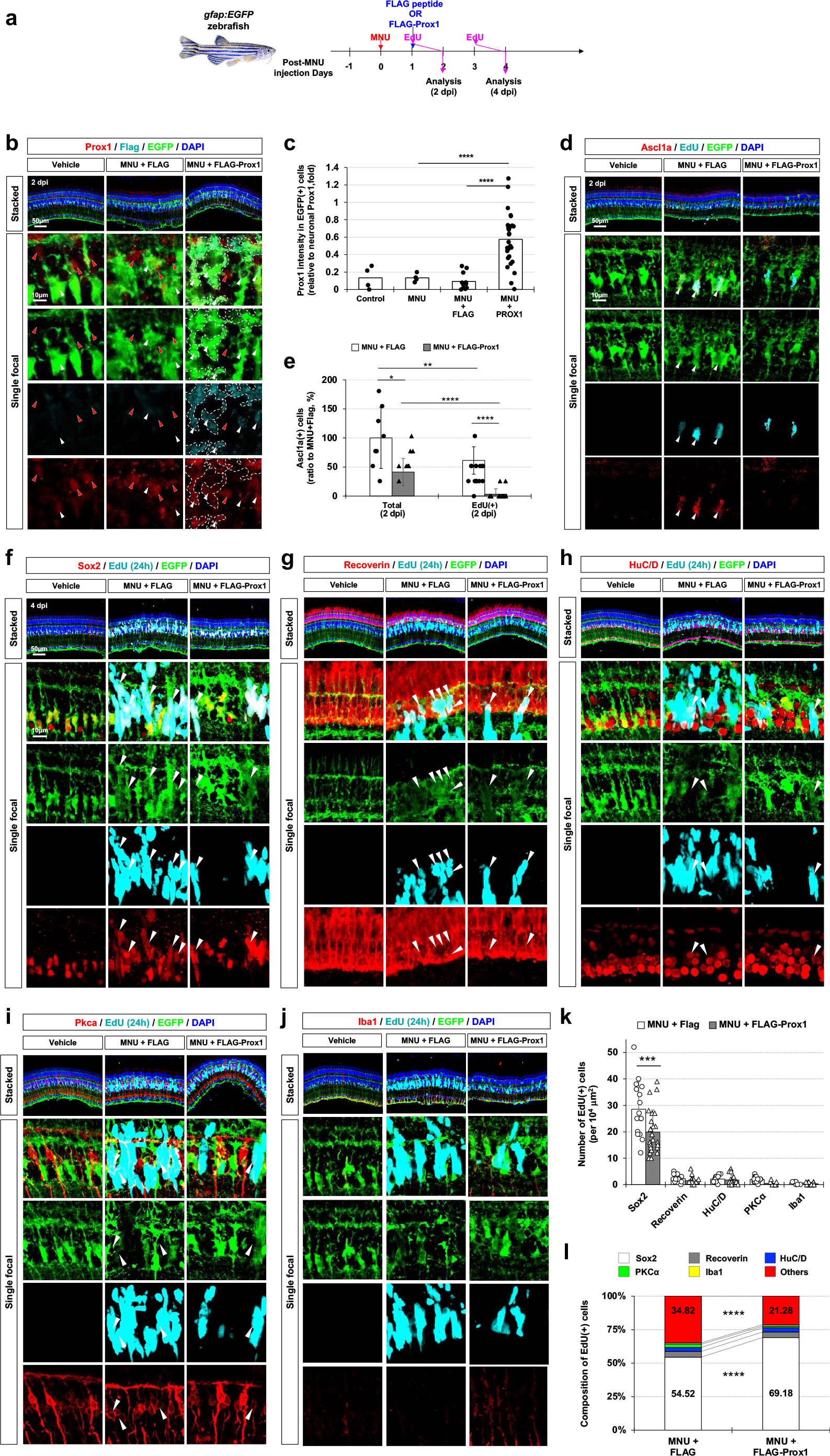
Exogenous Prox1 suppresses MG proliferation in the injured zebrafish retina. (a) FLAG peptides or recombinant FLAG-Prox1 proteins were injected into the intravitreal space of MNU-injured *gfap-EGFP* zebrafish eyes, and proliferating cells were labeled with EdU as it is indicated. **(b)** Prox1 distribution in the zebrafish retinas was examined by immunostaining with a rabbit anti-Prox1 antibody, and FLAG-Prox1 was detected by immunostaining with a mouse anti-FLAG antibody. White arrowheads indicate Prox1 in EGFP-positive MG, while red arrowheads point Prox1 in EGFP-negative retinal cells. The FLAG immunostaining signals are outlined by dotted lines. **(c)** The graph illustrates Prox1 intensity in MG relative to other retinal neurons within the same image (data from 3 independent litters). **(d)** MGPC identities of EdU-labeled cells in the retinas were determined by co-staining with the cell type-specific marker, Ascl1. White arrowheads point Ascl1;EdU-positive cells. **(e)** The number of Ascl1-positive cells in the zebrafish retinas is presented (data from 3 independent litters). **(f – j)** Additionally, the identities of EdU-labeled newborn cells in the zebrafish retinas were determined by co-staining with retinal cell type-specific markers, including Sox2 for MG or MGPCs (f); Recoverin for PR (g); HuC/D for AC (h); Pkcα for BC (i); and Iba1 for microglia (j). Arrowheads indicate EdU-labeled marker-positive cells. **(k)** The number of EdU-labeled cells expressing the markers are shown in the graph (data from 3 independent litters). **(l)** Cell composition of EdU-labeled cells in MNU-injured fish retinas is depicted in the graph. *p < 0.05; ** p < 0.01; *** p < 0.005 (ANOVA test).

**Extended Data Fig. 8.**
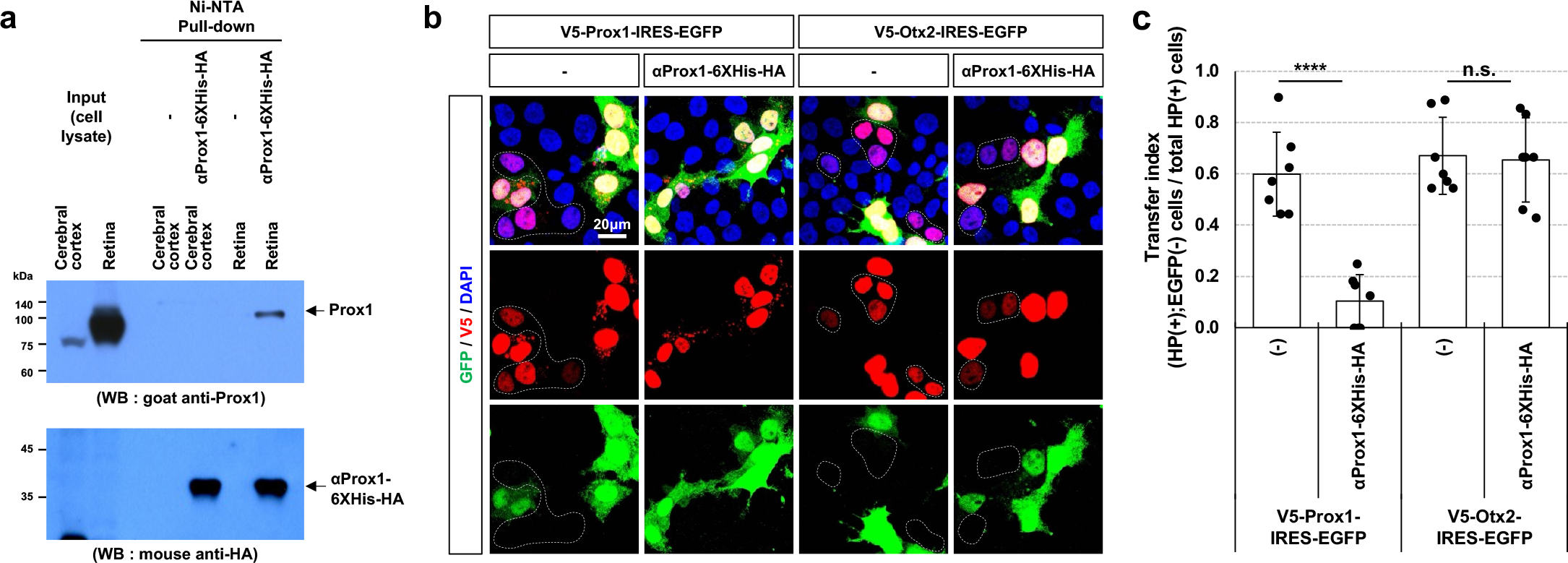
Identification of the neutralizing activity of αProx1 towards intercellular Prox1 transfer. (a) The cDNA of anti-Prox1 single-chain variable fragment (αProx1) was cloned into the M13 phagemid to express αProx1-6His-HA in Escherichia coli (E. coli), and the protein was purified using a Ni-NTA column, which captured its 6His tag of the antibody protein (see details in Methods). The αProx1-6His-HA protein (1 μg) was added to cell lysates (containing 1 mg protein) from the adult mouse cerebral cortex or retina and pulled down using the Ni-NTA column. Prox1, bound and co-purified with αProx1-6His-HA, was then detected by Western blot (WB) with goat anti-Prox1 antibody. The αProx1-6His-HA proteins in each lane were also detected by WB with mouse anti-HA antibody. **(b)** The αProx1-6His-HA protein was added to the growth medium of HeLa cells (25 μg/ml), which were transfected with pCAGIG-V5-Prox1 or pCAGIG-V5-Otx2. The cells expressing V5-Prox1 or V5-Otx2 with or without co-transcribed EGFP were identified by co-immunostaining with mouse anti-V5 and chicken anti-GFP antibodies. The cells surrounded by dotted lines contain the homeoproteins (HPs; i.e., Prox1 and Otx2) without EGFP, suggesting that they received the HP from HP;EGFP-positive neighboring cells. **(c)** The transfer index, which is the ratio of HP(+);EGFP(-) cells among total HP(+) cells, is depicted in the graph (data from 6 independent experimental batches).

**Extended Data Fig. 9.**
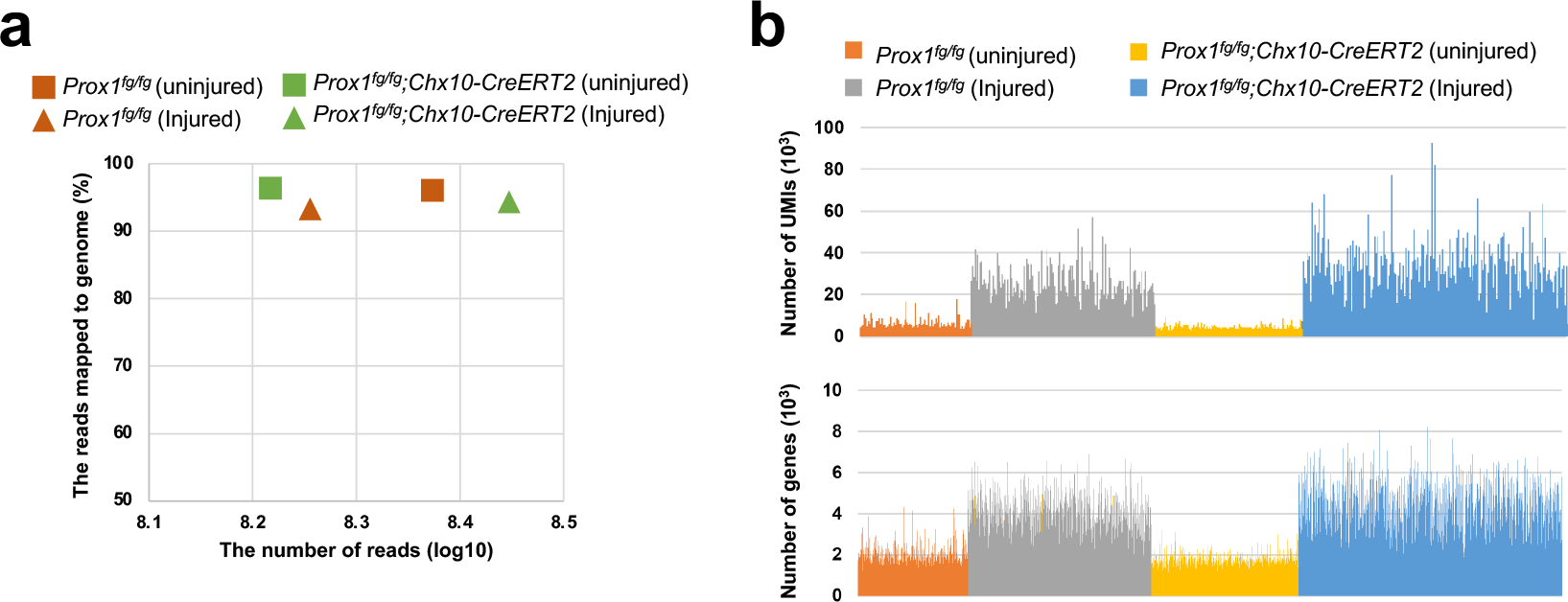
Quality of single-cell RNA-seq data. (a) Average number of reads from mouse retina cells with the indicated genotypes and those mapped to genome are presented in the graph. **(b)** Unique molecular identifiers (UMIs) and number of genes in each cell are depicted in the graphs.

**Extended Data Fig. 10.**
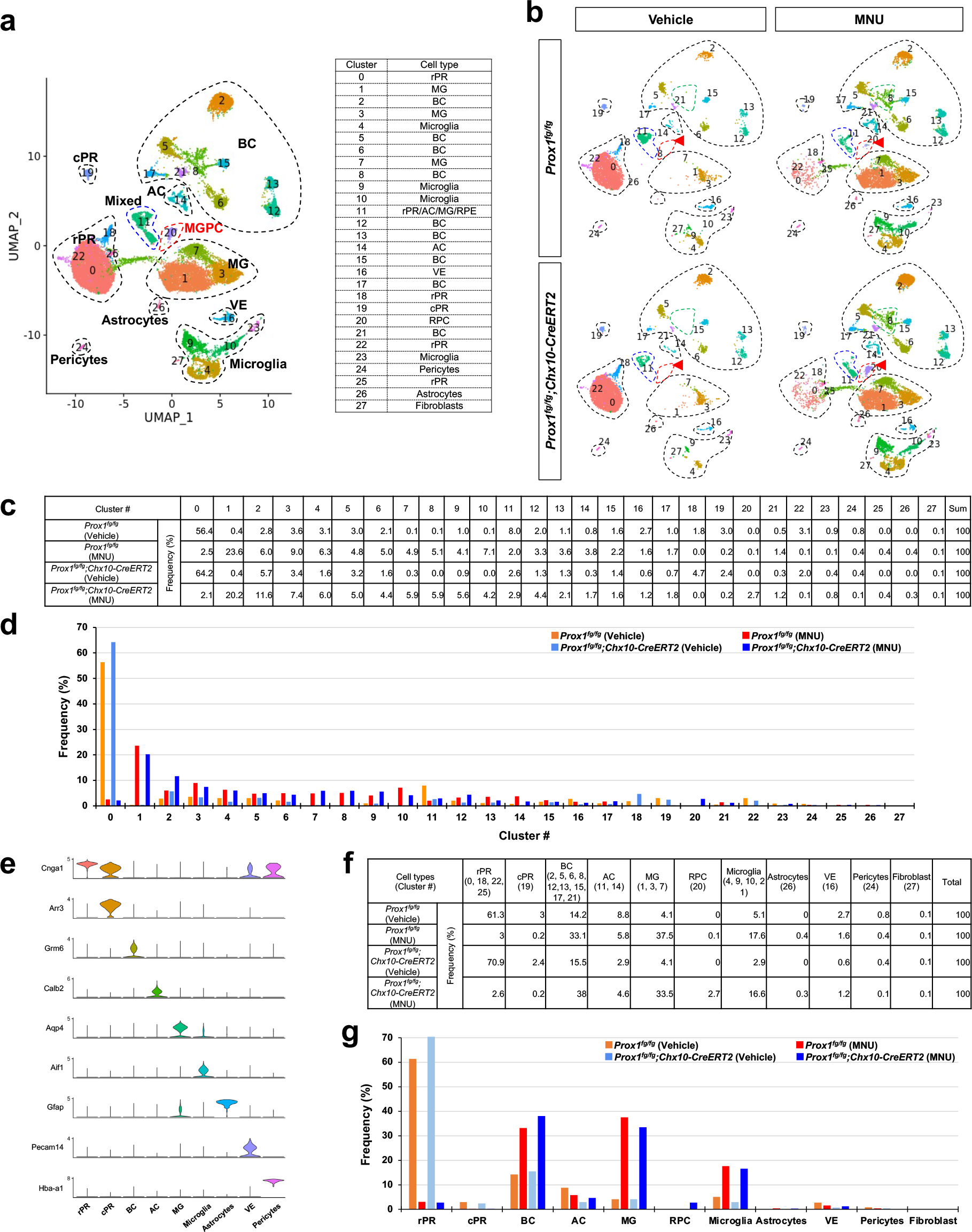
scRNA-seq analysis of the cells in MNU-injured mouse retinas. (a) UMAP of retinal cells, which were clustered by cell type. The identity of each cluster is provided in the table at the right. **(b)** Distribution of the cells in *Prox1^fg/fg^* and *Prox1^fg/fg^;Chx10-CreERT2* mouse retinas before (uninjured) and after (injured) MNU injury. Red arrowheads point cluster #20. **(c** and **d)** Frequency of each cluster in the mouse retinal cell population. (**e**) Violin plots of the expression of cell type-specific markers identified in previous studies. The cell type of each cluster was defined by these cell type-specific markers. **(f** and **g)** Frequency of each cell type in the mouse retinal cell population.

**Extended Data Fig. 11.**
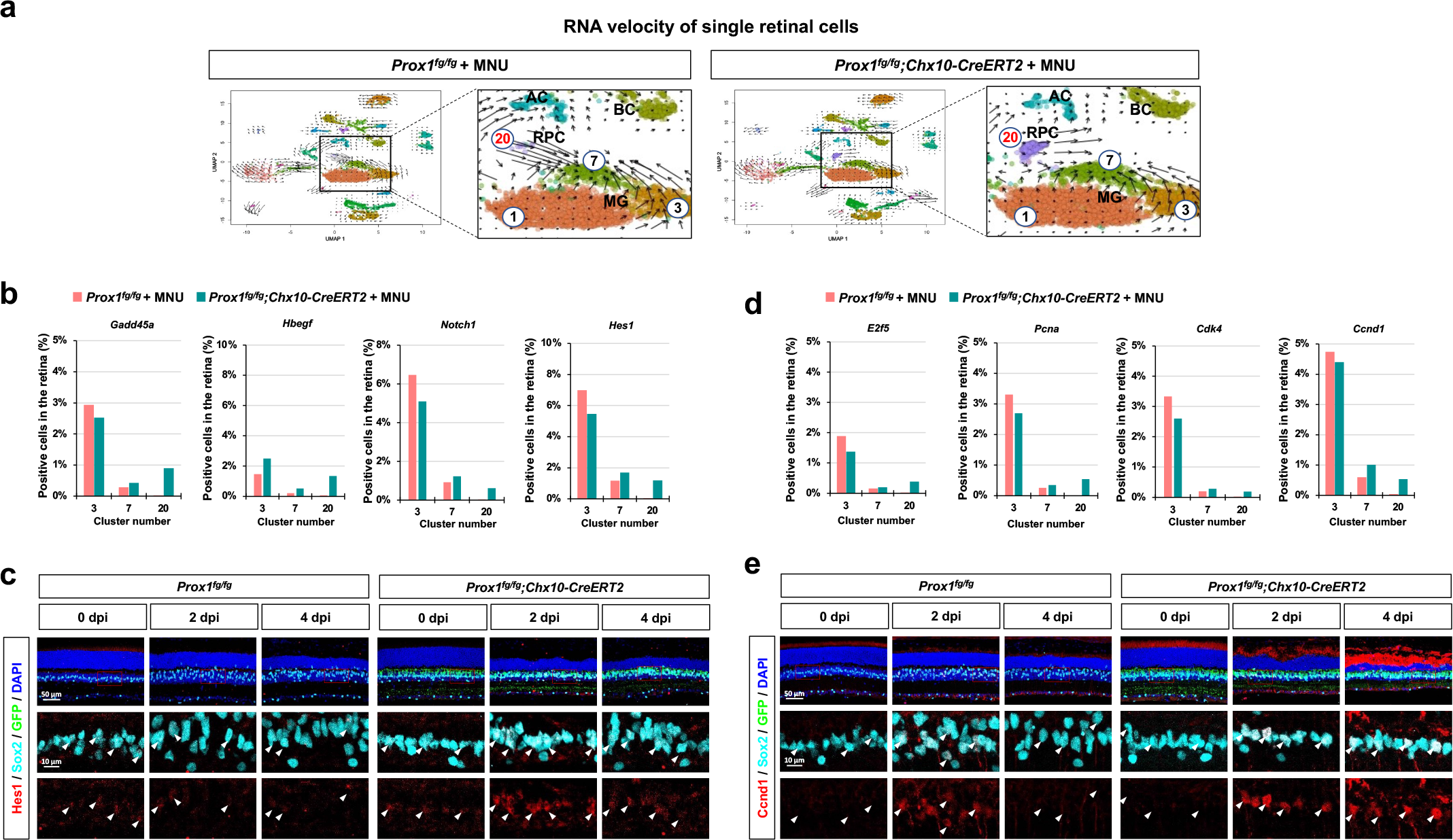
Injury-induced expression of Hes1 and Ccnd1 in MGPCs of *Prox1^fg/fg^;Chx10-CreERT2* mouse retinas. (a) RNA velocity analysis profiles of scRNA-seq results for injured *Prox1^fg/fg^* and *Prox1^fg/fg^;Chx10-CreERT2* mouse retinas. Images in the right are magnified versions of the boxed areas in the left. **(b)** The plots show the frequencies of cells expressing *Gadd45a*, *Hbegf*, *Notch1*, and *Hes1*, of which violin plots are provided in Fig. 3a. **(c)** Expression of Hes1 in Sox2-positive MG or MGPCs in the injured *Prox1^fg/fg^* and *Prox1^fg/fg^;Chx10- CreERT2* mouse retinas mouse retinas was investigated by immunostaining. **(d)** The plots show the frequencies of cells expressing *E2f5*, *Pcna*, *Cdk4*, and *Ccnd1*, of which violin plots are provided in Fig. 3d. **(e)** Expression of Ccnd1 in Sox2-positive MG or MGPCs in the injured *Prox1^fg/fg^*and *Prox1^fg/fg^;Chx10-CreERT2* mouse retinas mouse retinas was investigated by immunostaining. Arrowheads in (c) and (e) indicate Sox2-positive MG or MGPC nuclei.

**Extended Data Fig. 12.**
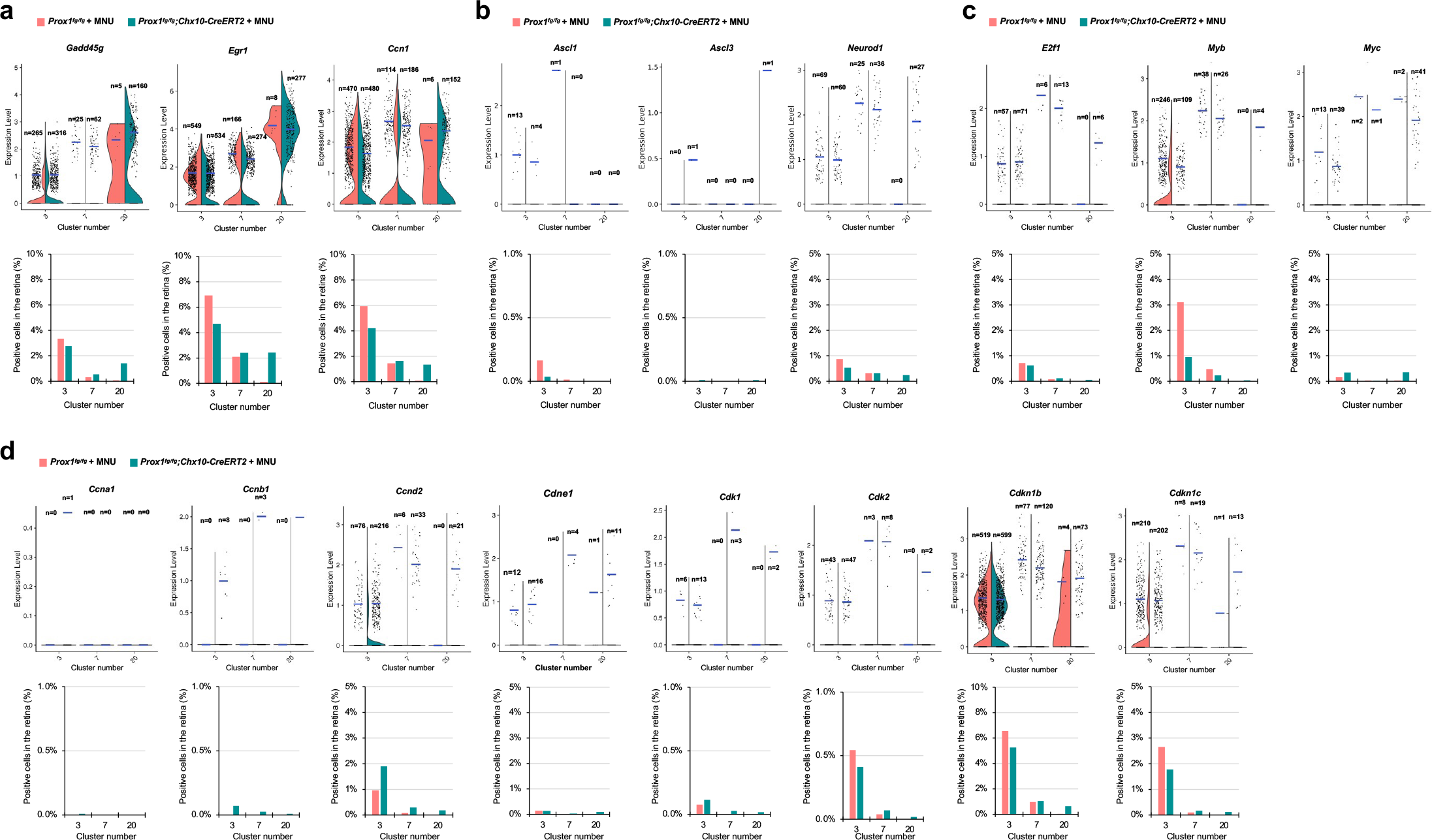
Expression of cell cycle regulatory genes in cluster #20. (a) Violin plots of the cells expressing mRNA of the RPC marker, *Gadd45g*; the target of Hbegf signaling, *Egr1*; and the target of Yap/Taz transcription activators, *Ccn1*, in MG (#3 and #7) and MGPC (#20) populations. **(b)** Violin plots of the cells expressing proneural transcription factor *Ascl1*, *Ascl3*, and *Neurod1* mRNA. **(c)** Violin plots of the cells expressing *E2f1*, *Myb*, and *Myc* mRNA. **(d)** Violin plots of the cells expressing Cyclin (*Ccna1*, *Ccnb1*, *Ccnd2*, and *Ccne1*), Cdk (*Cdk1* and *Cdk2*), and Cdk inhibitor (*Cdkn1b* and *Cdkn1c*) mRNA. Each dot represents the expression level of the corresponding gene in a cell. Blue horizontal bars indicate the mean expression. Graphs in the bottom of the plots show the percentage of the mRNA-expressing cells in total retinal cells.

**Extended Data Fig. 13.**
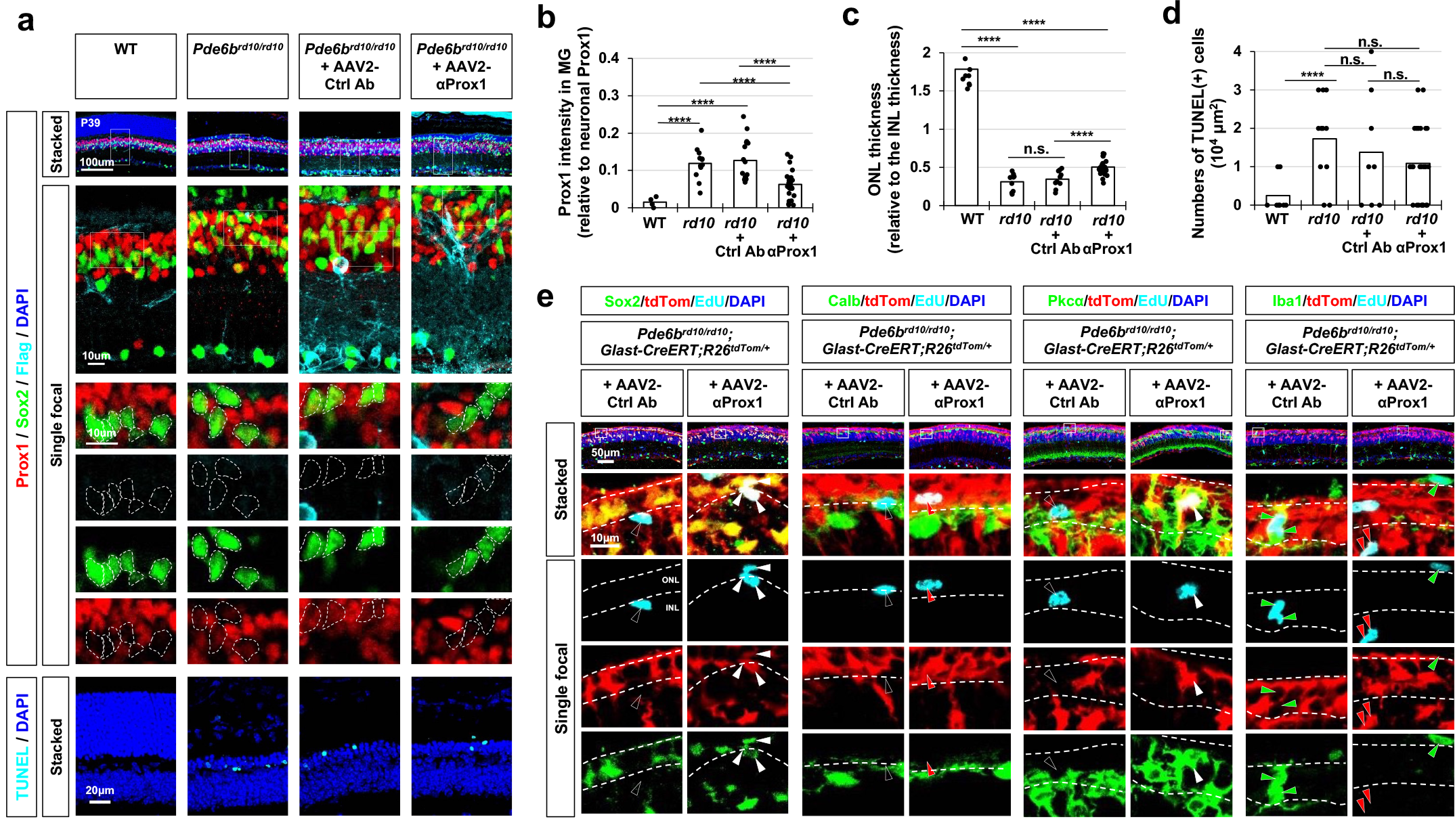
Regeneration of retinal neurons from MG in *Pde6b^rd10/rd10^* mice expressing αProx1. (a) *Pde6b^rd10/rd10^* mice were intravitreally injected with AAV2 encoding FLAG-tagged control antibody (Ctrl Ab) or αProx1, as shown in Fig. 4f. The expression of the antibodies in the mouse retinas was identified by immunostaining with mouse anti-FLAG antibody. Prox1 expression in Sox2-positive MG or MGPCs (outlined by dotted lines) were also examined by co-immunostaining with rabbit anti-Prox1 and goat anti-Sox2 antibodies. The apoptotic cells in the sections were identified by TUNEL staining. **(b)** Prox1 intensity in Sox2- positive MG or MGPCs relative to other retinal neurons in the same image is presented in the graph (data from 3 independent litters). **(c)** The thicknesses of the ONL and INL of the retinal sections were measured and relative thickness of the ONL compared to the INL of the same retinal sections is presented in the graph. **(d)** The number of TUNEL(+) apoptotic cells per retinal area is presented in the graph. **(e)** Identities of EdU-labeled cells in the AAV-injected *Pde6b^rd10/rd10^*mouse retinas were investigated by immunostaining for Sox2 (MG), Calb (HC), Pkcα (BC), and Iba1 (Microglia). MG origin of the cells was determined by the expression of tdTom Cre reporter. White arrowheads, EdU-positive cells co-expressing tdTom and the markers; red arrowheads, EdU-labeled cells containing tdTom but lacking of the markers; green arrowheads, EdU-labeled cells containing the markers but lacking of tdTom; black arrowheads, EdU-labeled cells lacking of tdTom and EdU. Numbers of EdU-positive cells expressing tdTom and corresponding markers are shown in Fig. 4h.

**Extended Data Fig. 14.**
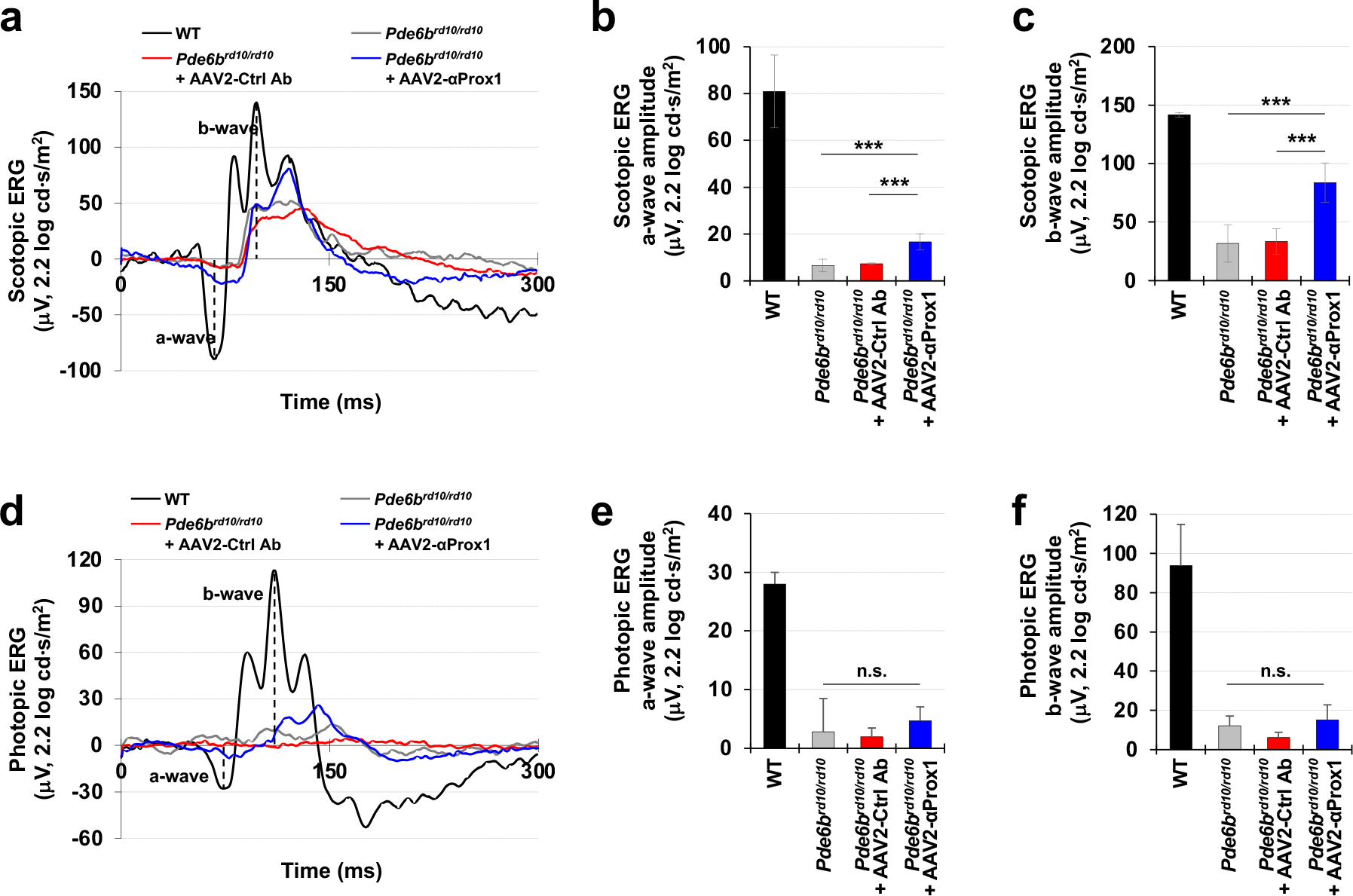
Recovery of photoreceptor activity in *Pde6b^rd10/rd10^* mice expressing αProx1. **(a** and **d)** Scotopic and photopic ERG histograms of *Pde6b^rd10/rd10^* mice infected with the indicated AAV2 were measured at P35 (see details in Methods). Amplitudes of ERG a-waves **(b** and **e)** and b-waves **(c** and **f)** in the mouse retina. Error bars in the graphs denote SEM (n=8; 4 independent litters). *** p<0.005 (ANOVA).

**Extended Data Fig. 15.**
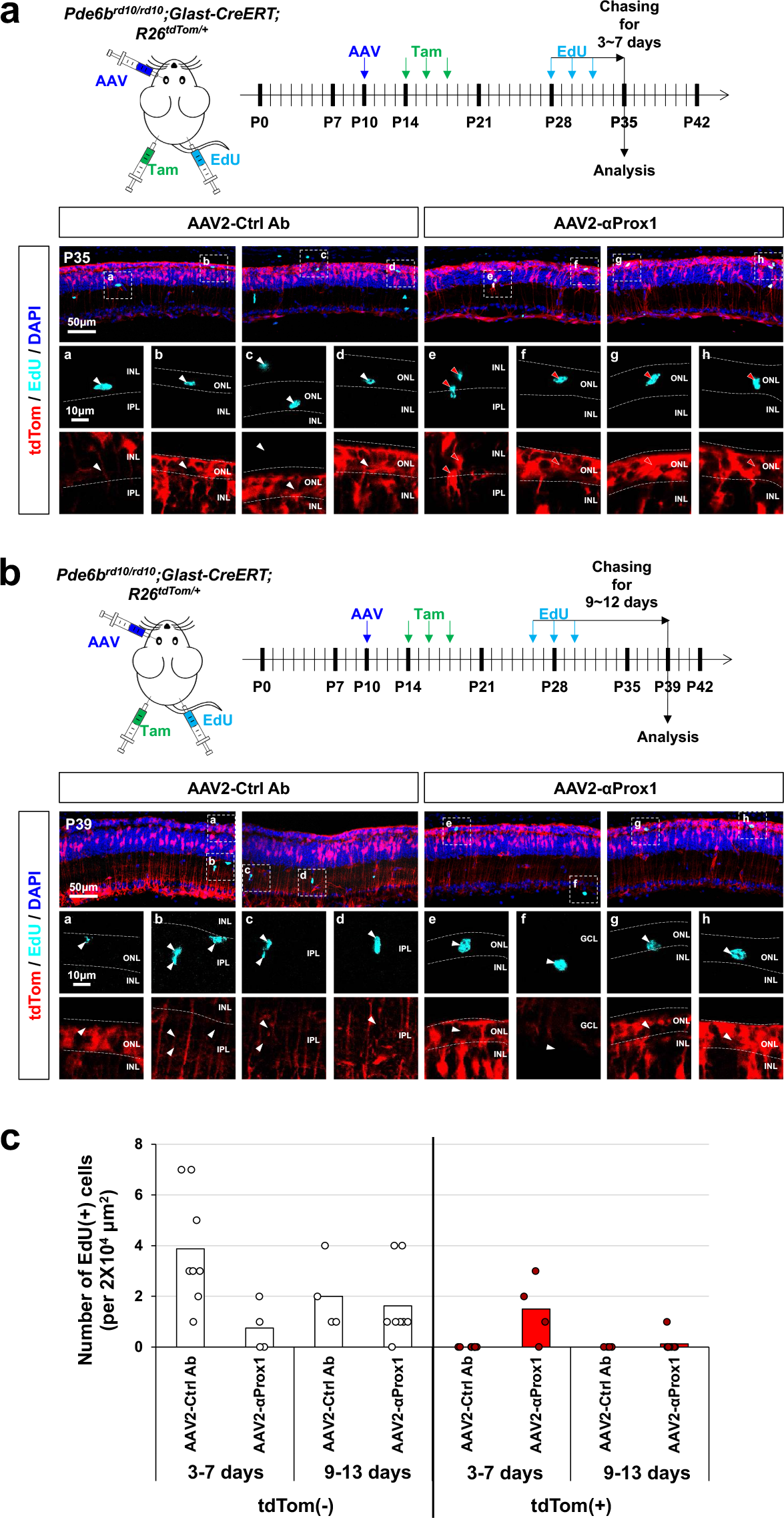
Lifespan of newborn photoreceptors in *Pde6b^rd10/rd10^* mouse retinas. **(a** and **b)** *Pde6b^rd10/rd10^;Glast-CreERT;R26^tdTom/+^*mice were intravitreally injected with AAV2 encoding FLAG-tagged Ctrl Ab or αProx1. The mice were repeatedly injected with Tam to activate CreERT in MG and EdU to labeled newborn cells in the retina as it is indicated. The mouse eyes were then isolated for the immunostaining of EdU-labeled cells in the retinal sections. MG origin of EdU-labeled cells was determined by the expression of tdTom Cre reporter. **(c)** Numbers of EdU-labeled cells remained by 3-7 days (a) or 9-12 days (b) in *Pde6b^rd10/rd10^*mouse retinas are presented in the graph (data from 3 independent litters).

**Extended Data Fig. 16.**
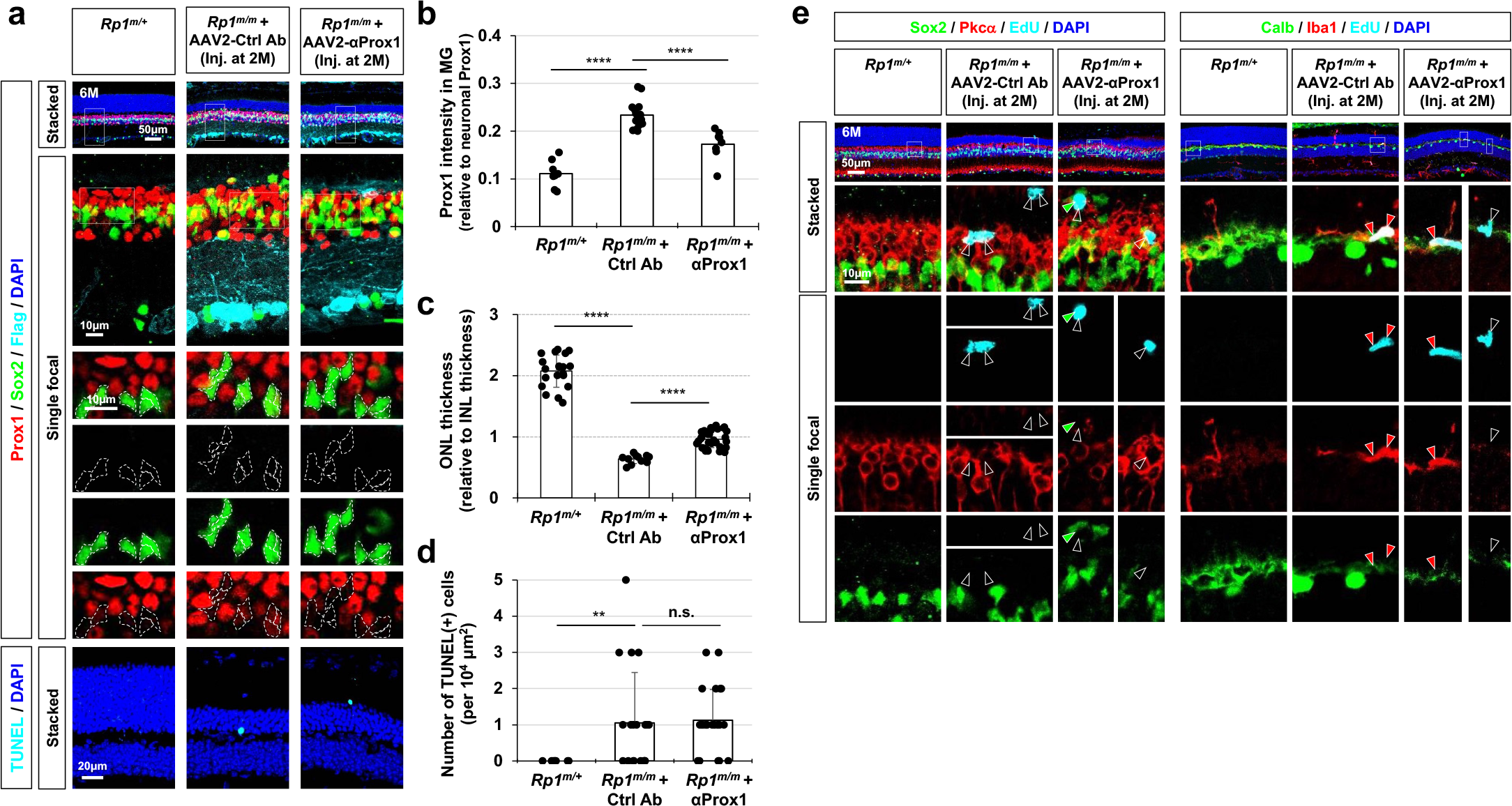
Regeneration of retinal neurons in *Rp1^m/m^* mice expressing αProx1. **(a)** *Rp1^m/m^* mice were intravitreally injected with AAV2 encoding FLAG-tagged Ctrl Ab or αProx1 at 2M. The expression of the antibodies in the mouse retinas was then identified at 6M by immunostaining with mouse anti-FLAG antibody. Prox1 expression in Sox2-positive MG or MGPC (surrounded by dotted circles) were examined by co-immunostaining with rabbit anti-Prox1 and goat anti-Sox2 antibodies. The apoptotic cells in the sections were identified by TUNEL staining. **(b)** Prox1 intensity in Sox2-positive MG relative to other retinal neurons in the same image is presented in the graph (data from 2 independent litters). **(c)** The thicknesses of the ONL and INL of the retinal sections were measured and relative thickness of the ONL compared to the INL of the same retinal sections is presented in the graph. **(d)** The number of TUNEL(+) apoptotic cells per retinal area is presented in the graph. **(e)** Identities of EdU-labeled cells in the AAV-injected *Rp1^m/m^* mouse retinas were investigated by immunostaining for Sox2 (MG or MGPC), Pkcα (BC), Calb (HC), and Iba1 (microglia). Red and green arrowheads, EdU-labeled cells containing the markers in corresponding colors; black arrowheads, EdU-labeled cells lacking of the markers. Numbers of EdU-positive cells expressing corresponding markers are shown in Fig. 5d. ****, p<0.001; **, p<0.01; n.s., not significant (data from 2 independent litters).

**Extended Data Fig. 17.**
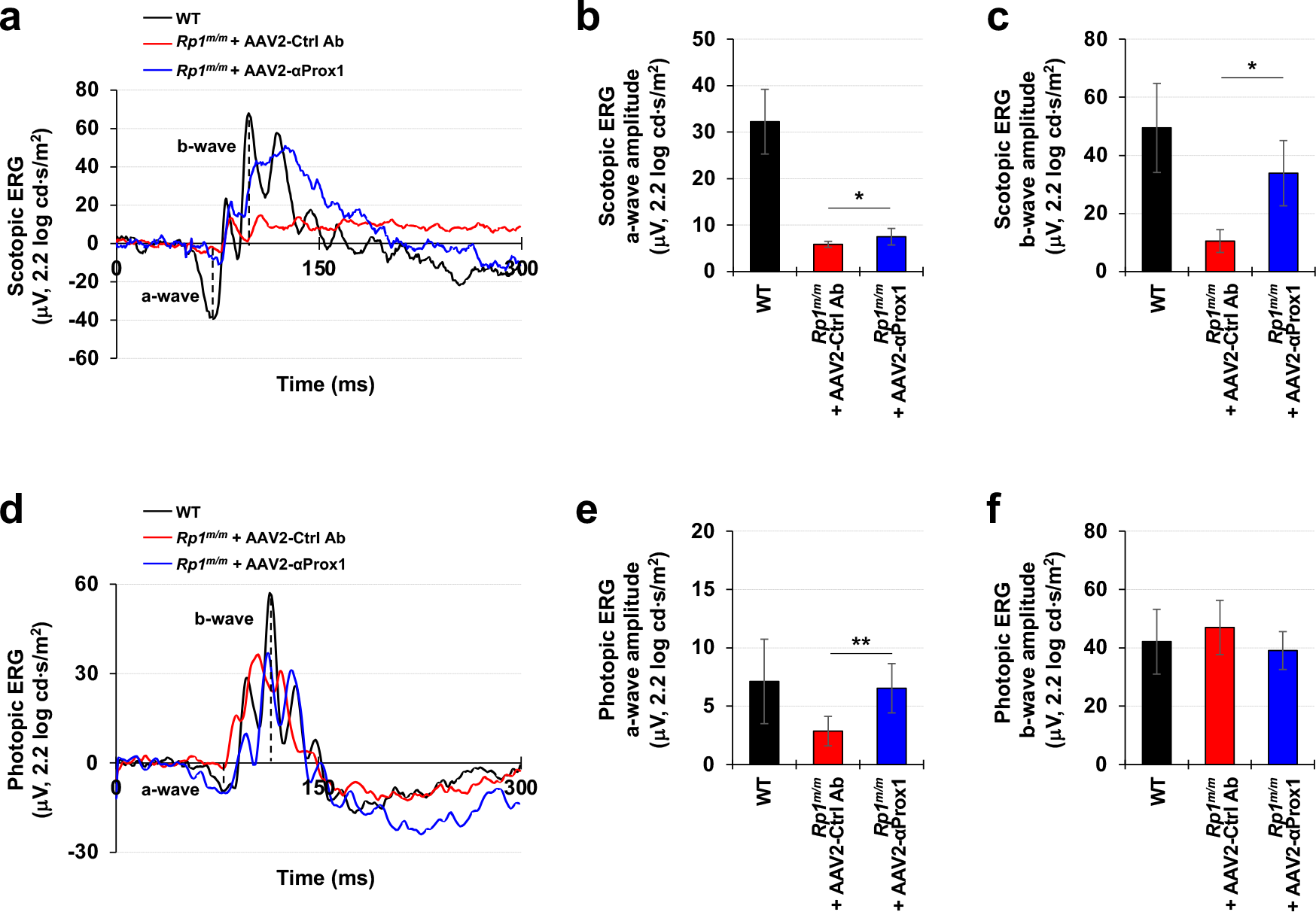
Recovery of photoreceptor activity in RP model mice expressing αProx1. **(a** and **d)** Scotopic and photopic ERG histograms of 6 months-old *Rp1^m/m^* mice infected with the indicated AAV2 were measured. Amplitudes of ERG a-waves **(b** and **e)** and b-waves **(c** and **f)** in the mouse retina. Error bars in the graphs denote SEM (n=6; 2 independent litters). * p<0.05; ** p<0.01 (ANOVA).

**Extended Data Fig. 18.**
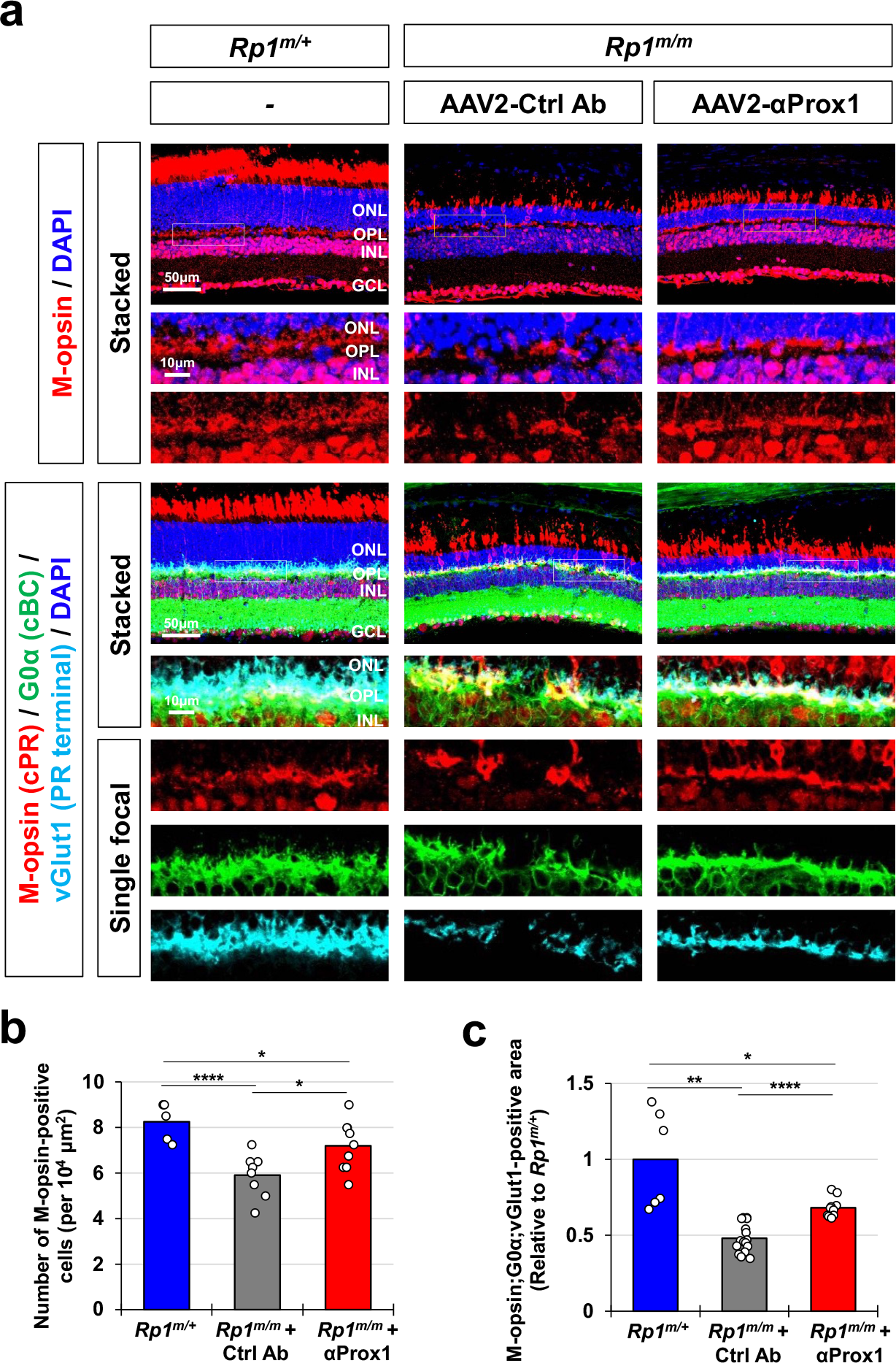
Preservation of cone photoreceptors in *Rp1^m/m^* mice expressing αProx1. (a) *Rp1^m/m^* mice were intravitreally injected with AAV2 encoding FLAG-tagged Ctrl Ab or αProx1, as shown in Fig. 5c Distribution of cPRs in the mouse retinas were examined by immunostaining of M-opsin. Synaptic terminals of the PRs and their post-synaptic targets, i.e., cone bipolar cells (cBCs), are visualized by co-immunostaining of respective markers, vGlut1 and G0α. **(b)** Number of M-opsin-positive cPRs were counted and presented in the graph (data from 2 independent litters). **(c)** The areas commonly exhibiting M-opsin;G0α;vGlut1 signals were measured from the immunostaining images and depict in the graph. * p<0.05; ** p<0.01; **** p<0.001.

**Supplementary table 1.**
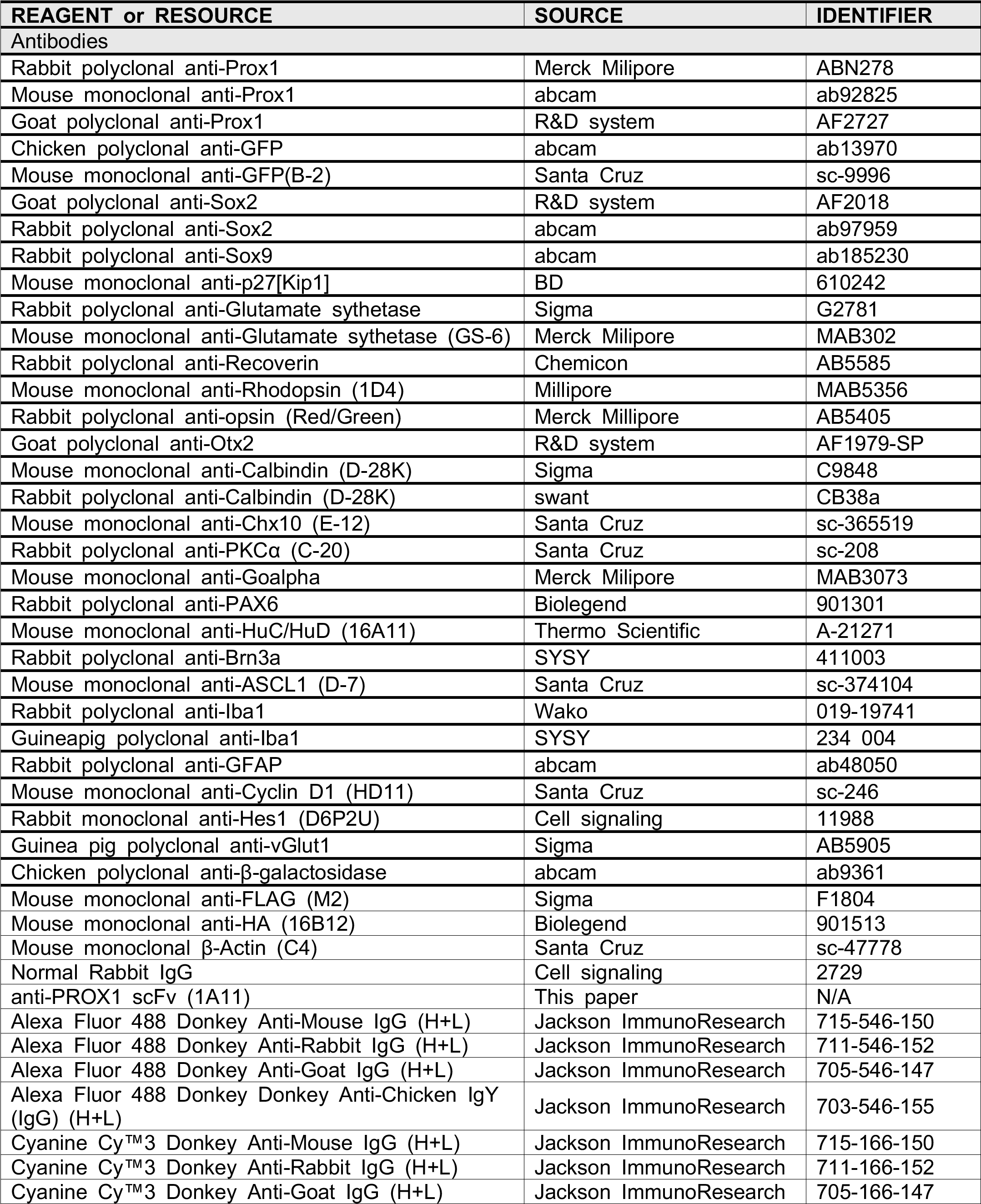
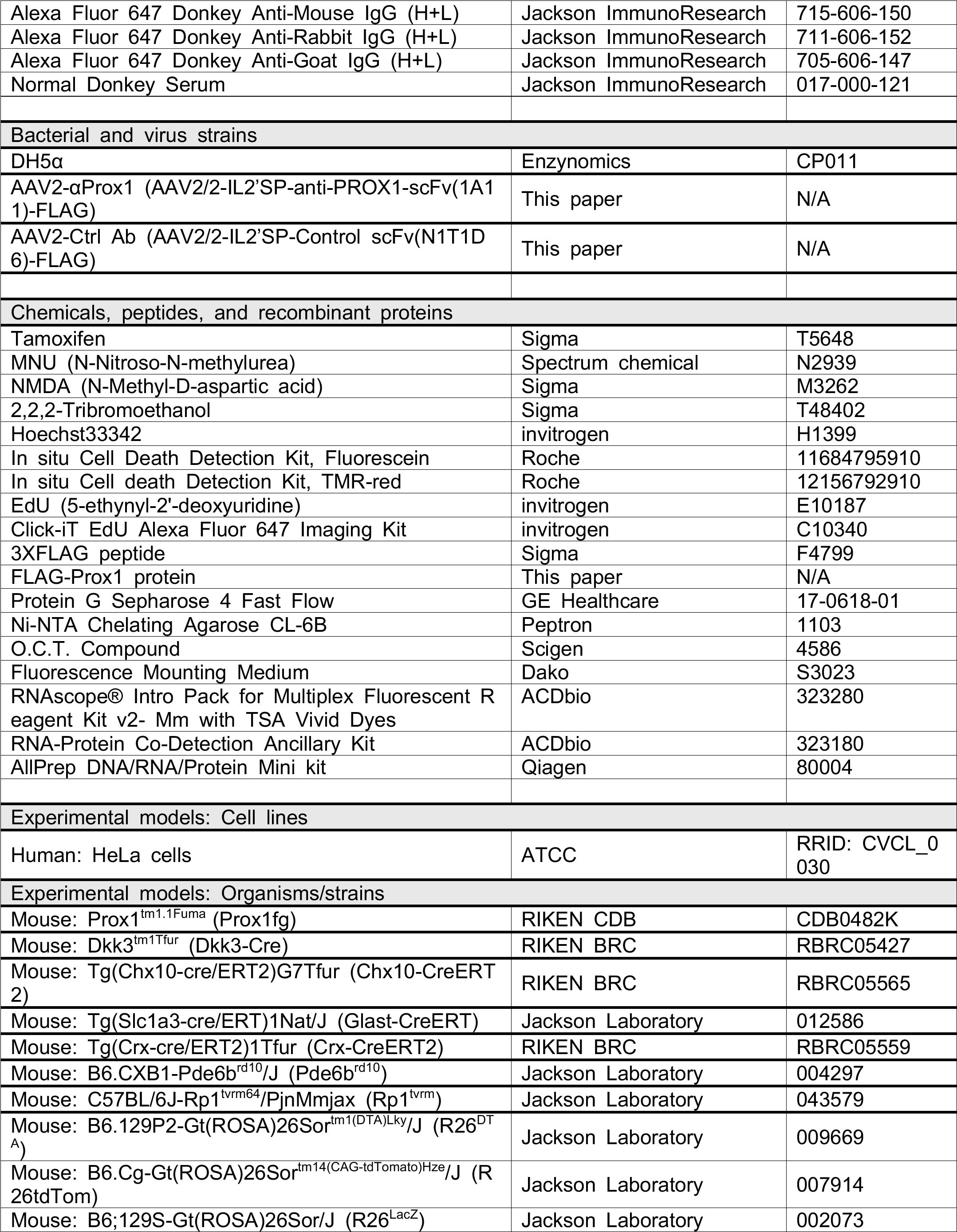
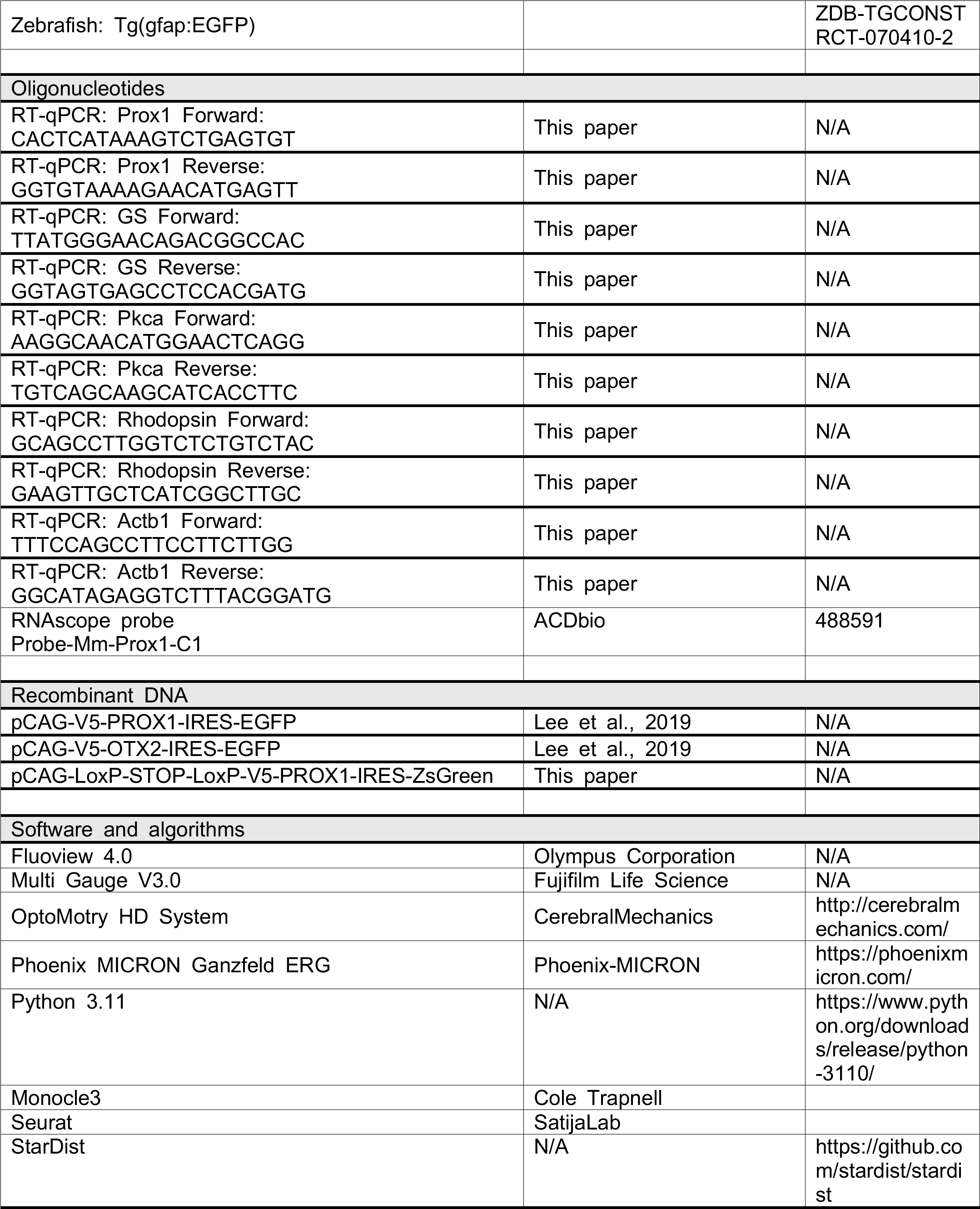
List of materials used in this study.

